# Human TRMT2A methylates tRNA and contributes to translation fidelity

**DOI:** 10.1101/2022.12.28.522094

**Authors:** Monika Witzenberger, Sandra Burczyk, David Settele, Wieland Mayer, Luisa M. Welp, Matthias Heiss, Mirko Wagner, Thomas Monecke, Robert Janowski, Thomas Carell, Henning Urlaub, Stefanie M. Hauck, Aaron Voigt, Dierk Niessing

## Abstract

Methyl-5-uridine (m5U) is one of the most abundant RNA modifications found in cytosolic tRNA. tRNA methyltransferase 2 homolog A (hTRMT2A) is the dedicated mammalian enzyme of m5U formation at tRNA position 54. However, its RNA binding specificity and functional role in the cell are not well understood. Here we dissected structural and sequence requirements for binding and methylation of its RNA targets. Specificity of tRNA modification by TRMT2A is achieved by a combination of modest binding preference and presence of a uridine in position 54 of tRNAs. Mutational analysis together with crosslinking experiments identified a large hTRMT2A-tRNA binding surface. Furthermore, complementing hTRMT2A interactome studies revealed that TRMT2A interacts with proteins involved in RNA biogenesis. Finally, we addressed the question of the importance of TRMT2A function by showing that its knockdown reduces translation fidelity. These findings extend the role of hTRMT2A beyond tRNA modification towards a role in translation.

## Introduction

Up to date more than 170 post-transcriptional modification were reported. They can affect RNA stability, translation, and localization and, hence, constitute an important layer of gene regulation (1–3). Amongst the different types of transcripts, tRNA is the most heavily modified RNA species with an average of 13 modifications per tRNA molecule (4–6). These modifications play a role in tRNA stability and folding, although their exact biological function often remains elusive. The modification 5-methyluridine (m^5^U) frequently occurs at position 54 in the T-Loop of bacterial and eukaryotic tRNAs (4). While in *E. coli* the tRNA (uracil-C(5))- methyltransferase A (TrmA) is responsible for m^5^U formation, the respective homolog in *S. cerevisiae* is termed Trm2, and in mammals tRNA (uracil-5)-methyltransferase homolog A (TRMT2A) (7–10). In addition, mammals possess TRMT2B, the mitochondria-specific paralog (11, 12).

Removal of the catalytic activity of TrmA does not impact the viability of *E. coli*, although deletion of the whole gene is lethal (13). In contrast, in budding yeast Trm2 is not essential for survival (9). Hence, these phenotypes suggest a supporting role of the enzyme in tRNA maturation and other yet unknown biological processes beyond m^5^U catalysis (14). For instance, *E. coli* TrmA has recently been reported to be a tRNA chaperone assisting proper tRNA folding (15). Moreover, TrmA methylates not only tRNA, but also the T-Loop of tmRNA (16) and 16S rRNA *in vitro* (17).

Detailed biochemical and structural studies on *E. coli* TrmA have contributed to a better understanding of the general enzymatic mechanism for m^5^U formation. Most methyltransferases, including TrmA as well as human TRMT2A, use SAM (S-adenosyl- methionine) as methyl donor (10, 12, 18). Previous mutational studies identified catalytically relevant residues in TrmA. Enzymatic activity as well as RNA binding was reduced upon mutation of the catalytic cysteine (C324A). The putative SAM binding site (G220D), the proton abstraction base for the catalytic reaction (E358K) and one of the residues interacting with target uridine (Q190A) were important for RNA binding too (18). Furthermore, residues (H125A, F106) reduced tRNA binding and methylation, since they disrupt the tertiary interaction of T- and D-Loop (15). Such biochemical studies were supported by a co-crystal structure of TrmA with the T-Loop of tRNA (19), confirming that the conserved C324 covalently binds to the target uridine, thus driving m^5^U formation.

Only recently our knowledge about m^5^U tRNA methyltransferases was expanded to human TRMT2A (hTRMT2A). Transcriptome-wide mapping of hTRMT2A methylation sites on RNA using FICC-CLIP have revealed that 87 % of the cellular hTRMT2A targets are indeed tRNAs (10). Binding sequences with the motif CTTCG/AA were enriched, which constitutes a conserved tRNA T-Loop sequence. However, hTRMT2A binding expands beyond tRNAs as observed by its interaction with the HIST1H4B or KCND2 mRNA, small subunit rRNA and other RNAs (20, 21). In line with such a broader function, its mammalian paralog TRMT2B was not only shown to be the dedicated mitochondrial methyltransferase for m^5^U54 formation in mitochondrial tRNAs, but also to promote m^5^U429 formation of mitochondrial 12S rRNA (11, 12). It remains to be shown whether also hTRMT2A has a dual function like methylating tRNA and rRNA.

The cellular role of hTRMT2A, m^5^U formation and possible roles beyond m^5^U formation are not well understood, although some functional observations have been made; while in one study, hTRMT2A overexpression correlated with Her^+^ positive cancer (22) in another study hTRMT2A was described as cell cycle regulator that suppresses cell proliferation (23). In a third study, hTRMT2A KO caused accumulation of tRNA-derived fragments (tRFs) (24). Although the hTRMT2A-bound RNAs have been identified (10), we lack an understanding of its protein interactome and hence an overview of the molecular pathways it contributes to. Furthermore, to date all molecular properties of hTRMT2A have been inferred from orthologs in single-cell organisms.

In this study, we systematically characterized the functional and molecular features of hTRMT2A. Moreover, we identified hTRMT2A’s determinants of specificity. We observed that hTRMT2A has only a modest binding preference for its physiological targets and that target specificity is rather achieved by the requirement of a uridine in the correct steric position to allow for site-specific methylation. Assessment of the cellular human TRMT2A interactome allowed us to identify potential co-factors, suggesting novel roles of hTRMT2A in different cellular processes. Finally, we show that loss of hTRMT2A in a cellular context resulted in reduced translation fidelity.

## Materials and Methods

### Cloning, expression in *E. coli* and purification

Plasmid containing human TRMT2A sequence was provided by Aaron Voigt. Human TRMT2A RBD (aa 69 – 147), Central domain (aa 237-414) and Fusion domain (aa 68-240/413-590) were cloned with a cleavable SUMO-tag at the N-terminus and expressed in Rosetta 2 (DE3) in case of RBD or Rosetta 2 (DE3) pLyS (Central, Fusion) *E. coli* in LB medium. After isopropyl-thiogalactoside (IPTG, 0.5 mM) induction overnight at 18 °C, cells were harvested (4500 x *g*, 15 min, 4 °C). All purification steps were performed at 4 °C. For hTRMT2A RBD purification typically pellets from 6 liters of culture were resuspended in lysis buffer-A (50 mM HEPES-NaOH pH 8.5, 500 mM NaCl, 20 mM Imidazole, 0.5 % (v/v) Tween, 2 % Glycerol) supplemented with one tablet of EDTA-free protease inhibitor cocktail (Roche). After sonication (4x 6 min, amplitude 40 %), the lysate was clarified by centrifugation (40,000 x *g*, 30 min). The lysate was loaded onto a 5 ml HisTrap FF column (GE Healthcare), equilibrated in His-A buffer (50 mM HEPES-NaOH pH 8.5, 500 mM NaCl, 20 mM Imidazole), and washed with 10 column volumes (CV) of His-A buffer, followed by a 10 CV wash with His-B buffer (50 mM HEPES-NaOH pH 8.5, 2000 mM NaCl, 20 mM Imidazole). SUMO-tagged protein was eluted with a 10 CV gradient of His-A buffer and His-C elution buffer (50 mM HEPES-NaOH pH 8.5, 500 mM NaCl, 500 mM Imidazole). For tag cleavage, the eluate was supplemented with 100 µg of PreScission™ protease and dialyzed against dialysis buffer-A (50 mM HEPES-NaOH pH 7.5, 500 mM NaCl, 1 mM DTT) overnight. Cleaved protein was run on a subtractive HisTrap FF column and the protein-containing flow-through was collected and concentrated. Protein was loaded onto a size exclusion chromatography column (Superdex 75, 10/300 gl) equilibrated in SEC-A buffer (50 mM HEPES-NaOH pH 7.5, 500 mM NaCl). Protein concentrations were determined by measurement of the A280. Aliquots were flash-frozen in liquid nitrogen and stored at -80 °C.

For Central and Fusion domain, lysis buffer-B was used (500 mM NaCl, 50 mM HEPES-NaOH pH 7.5, 0.1 % (v/v) Tween, 20 mM Imidazol, 10 % Glycerol). Moreover, all buffers contained HEPES-NaOH with pH 7.5 and for the final size exclusion chromatography a Superdex 200 column (16/600) and SEC- B buffer were used (50 mM HEPES-NaOH pH 7.5, 500 mM NaCl, 1 mM DTT). Fusion domain was otherwise purified like hTRMT2A RBD. The SUMO-tag of the Central domain was not cleaved due to subsequent protein degradation. Instead, after overnight dialysis in dialysis buffer-B (50 mM HEPES-NaOH pH 7.5, 150 mM NaCl, 1 mM DTT) the protein was loaded onto a HiTrap Heparin column equilibrated with dialysis buffer-B and eluted with Hep-B elution buffer (50 mM HEPES-NaOH pH 7.5, 1000 mM NaCl, 1 mM DTT). Protein-containing fractions were processed as described above. Pooled protein peak fractions were concentrated to 2-5 mg/mL with a centrifugal filter (Amicon), flash-frozen in liquid nitrogen and aliquots were stored at -80 °C. Primers for cloning and mutagenesis are listed in **Supplementary Table 1**

### Human TRMT2A FL cloning, expression in insect cells, and purification

The Bac-to-Bac baculovirus expression system (Invitrogen) was employed for hTRMT2A FL expression. N-terminal SUMO-tag fused hTRMT2A FL and its mutants were cloned into the pFastBac vector. For transfection using the FuGene HD transfection reagent 1.5 µg bacmid DNA and 3 µL transfection reagent were preincubated for 30 min in 200 µL Sf-900 medium at room temperature, then added dropwise to the Sf21 cells pre-seeded on 6-well plates with a density of 0.4*10^6^ cells/mL. Then cells were incubated for 4 days at 27.5 °C.

Supernatant of transfected cells from two 6-wells was used to infect 10 mL of Sf21 shaking cell culture at a density of 1.4*10^6^ cells/mL. After 3-4 days of incubation at 27.5 °C, insect cells were harvested by centrifugation for 10 min at 4 °C with 2000 rpm (Hettich centrifuge Rotina 420R). Virus-containing supernatant P1 was sterile-filtered using a 0.22 μm filter (Merck). To amplify the virus 1.5 mL P1 was added to 250 mL of Sf21 shaking cell culture at a density of 0.4*10^6^ cells/mL. After another 3-4 days of incubation at 27.5 °C, cells were harvested. The virus-containing supernatant P2 was sterile filtered and stored at 4 °C. Protein was expressed in Hi5 insect cells by adding 20 mL of P2 to 500 mL shaking culture at a density of 1*10^6^ cells/mL. Proteins were expressed in Hi5 insect cells for 2-3 days at 27.5 °C.

Cells were harvested by centrifugation (2000 x *g*, 15 min, 4 °C) and resuspended in lysis buffer-C (500 mM NaCl, 50 mM HEPES-NaOH pH 7.5, 0.1 % (v/v) Tween, 10 % Glycerol, 20 mM Imidazol). Next, insect cells were lysed using a dounce homogenizer. The lysate was clarified by centrifugation (40,000 x *g*, 30 min, 4 °C). Subsequent purification was performed as for hTRMT2A Fusion domain with an initial HisTrap FF column, cleavage of the SUMO tag that was followed by a subtractive HisTrap FF column, dialysis overnight with dialysis buffer-A and a final size exclusion chromatography run with a Superdex 200 column (16/600) in SEC-B buffer. hTRMT2A FL for crosslinking and mass spectrometry experiments was purified with SEC-C buffer (50 mM HEPES-NaOH pH 7.5, 500 mM NaCl, 0.5 mM TCEP), where DTT was exchanged by TCEP. Pooled protein peak fractions were concentrated to 2- 5 mg/mL with a centrifugal filter (Amicon), flash-frozen in liquid nitrogen and aliquots were stored at - 80 °C.

### RNA in vitro transcription and purification

All RNAs were transcribed from a HPLC-purified forward primer, which contained the T7 RNA polymerase promoter region and a PAGE-purified reverse primer that consisted of the reverse complement DNA template and the T7 RNA polymerase promoter region (**Supplementary Table 2**). Briefly, 4 µM forward primer and 3.3 µM reverse primer were annealed in the presence of 20 mM MgCl2 for 5 min at 60 °C in a total volume of 1.2 mL and subsequently cooled to room temperature. 4 mM of each NTP, 16 mM MgCl2, 80 mg/mL PEG 800, 0.5 mg/mL T7 RNA Polymerase and 10X TRX-buffer (40 mM Tris/Cl pH 8.1, 1 mM spermidine, 0.01 % Triton X-100, 5 mM DTT) were added to the annealed primer-pair in a final volume of 5 mL. After incubation for 3 h at 37 °C, the precipitate was removed by centrifugation for 10 min at 4 °C with 14,000 x g. Thereafter the RNA was precipitated with 0.1 volumes 3 M sodium acetate and 3.5 volumes ethanol for > 30 min at -20 °C. RNA pellets were resuspended in 0.5 mL DEPC-H2O and 0.5 mL denaturing loading dye was added (2 mM Tris/Cl pH 7.5, 20 mM EDTA, 8 M urea, 0.025 % (w/v) bromophenol blue, 0.025 % (w/v) xylene cyanole). Separation on 8 % denaturing TBE-PAGE was followed by visualization of transcripts by UV shadowing and excision from gel. RNA was extracted by electroelution (Whatman Elutrap, GE Healthcare) at 200 V in 1X TBE according to manufacturer’s instructions. Fractions containing the RNA were pooled dialyzed twice against 5 M NaCl and once against DEPC-H2O. RNAs were lyophilized, subsequently resuspended in DEPC-H2O and stored at - 20 °C. General recommendations for working with RNAs were followed (25).

### RNA radiolabeling and electromobility shift assays (EMSAs)

*In vitro* transcribed RNA was 5’ dephosphorylated using FastAP™ thermosensitive alkaline phosphatase (Thermo Scientific) according to manufacturer’s recommendations. Thereafter, RNA was purified with Phenol-chloroform-isoamyl- alcohol extraction followed by isopropanol precipitation. 13 pmol of purified RNA was phosphorylated with y-^32^P ATP (Hartmann Analytics) by T4 polynucleotide kinase (NEB) for 30 min at 37 °C. The enzyme was deactivated at 75 °C for 10 min and labeled RNA was purified with a NucAway Spin column (Ambion). To ensure proper RNA folding after each thawing cycle, RNA was denatured at 98 °C for 2 min and snap-cooled on ice prior to usage.

For 20 µL EMSA reactions increasing protein concentrations (0-25 µM) were incubated with 100 nM y-^32^P labeled RNA in the presence of 2.5 ng/µL polyU competitor (poly-uridine ssRNA, Sigma) for 20 min at room temperature in RNAse-free EMSA-buffer (50 mM HEPES-NaOH pH 7.5, 150 mM NaCl, 4 % (v/v) glycerol). This results in a final concentration of 5 nM labeled RNA and 0-1.25 µM protein per reaction. Complexes were separated on 4 % native TBE-PAGE in TBE running buffer at 80 V. After gel fixation with 30 % (v/v) methanol and 10 % (v/v) acetic acid solution for 10 min, gels were dried in a vacuum gel dryer (model 573, BioRad). For analysis phosphor imaging plates were exposed to the radioactive dried gels for > 20 min and scanned with a FLA-300 system (Fujifilm). EMSAs were performed in triplicates on different days.

### Methyltransferase Assay

hTRMT2A FL methylation activity was analyzed with the MTase-Glo™ assay (Promega). The assay was performed according to the manufacturer’s instructions and experiments described in (26) in white-opaque OptiPlate™-384 (PerkinElmer). Experiments to determine Km values were performed with increasing substrate concentrations. A final concentration of 5 μM hTRMT2A FL was incubated with final concentration of 0-6 µM *in vitro* transcribed tRNA and 20 µM SAM in 1X reaction buffer (20 mM Tris buffer pH 8.0, 50 mM NaCl, 1 mM EDTA, 3 mM MgCl2, 0.1 mg/mL BSA, 1 mM DTT) for 60 min at room temperature. The reaction was stopped by adding 0.5 % TFA (PanReac AppliChem). After 5 min incubation, 6X MTase-Glo reaction solution diluted in MilliQ- H2O was added and incubated for 30 min at room temperature. Finally, MTase-Glo detection solution was added, incubated for 30 min in the dark at room temperature, and luminescence was read out using a microplate reader (Perkin-Elmer). The data was transformed to display relative luminescence by subtracting the value of a negative control reaction without RNA. Data of triplicate measurements were plotted with R (version 4.1.0) and fitted with Michaelis Menten equation to obtain Km values.

### Crosslinking and Mass-spectrometry

For the first set of experiments complex reconstitution was performed; to do so *in vitro* transcribed tRNA^Gln^ and recombinantly purified hTMRT2A FL were reconstituted in reconstitution buffer (50 mM HEPES-NaOH pH 7.5, 50 mM NaCl, 0.5 mM TCEP) in a 1:4 ratio. For the second set of experiments, the complex was co-purified using size-exclusion chromatography. Briefly, tRNA^Gln^ and hTRMT2A FL were incubated in a 1:1 ratio on ice for 30 min. Then, the complex was separated from single RNA and protein using an S6 (15/150) column. For UV crosslinking, the *in vitro* reconstituted and co-purified complex was irradiated at 254 nm for 10 min on ice using an in-house built crosslinking apparatus as described previously (27). For chemical cross- linking, mechlorethamine (NM, B1785, APExBIO Technology LLC) was added to a final concentration of 1 mM following incubation for 30 min at 37 °C. Crosslinked samples were precipitated using ethanol and further processed as described in Kramer et al., 2014. In brief, 10 µg RNase A (EN0531, Thermo Fisher Scientific), 1 kU RNase T1 (EN0531, Thermo Fisher Scientific) and 250 U PierceTM universal nuclease (88700, Thermo Fisher Scientific) were used for tRNA digestion that was carried out for 2 h at 37 °C. Subsequently, proteins were digested using trypsin (V5111, Promega) at a 1:20 (enzyme to protein) mass ratio. After protein digestion, another 10 μg of RNase A and 1 kU of RNase T1 were added and the samples were incubated for 1 h at 37 °C. Free nucleotides and salts were removed using C18 columns (744601, Harvard Apparatus) and crosslinked peptides were enriched using in-house packed TiO2 columns (Titansphere 5 µm; GL Sciences). Enriched peptides were dried and dissolved in 2 % (v/v) acetonitrile, 0.05 % (v/v) TFA. LS-MS/MS analyses were performed on a Thermo Scientific Orbitrap Exploris 480 mass spectrometer coupled to a nanoflow liquid chromatography system (Thermo Scientific Dionex Ultimate 3000). Peptide separation was performed over 58 min with a flow rate of 300 nl/min using a linear gradient and a buffer system consisting of 0.1 % (v/v) formic acid (buffer A) and 80 % (v/v) acetonitrile, 0.08 % (v/v) formic acid (buffer B). Peptide-nucleotide hetero-conjugates were analysed in positive mode using a data-dependent top 30-acquisition method. MS1 and MS2 resolution were set to 120,000 and 30,000 FWHM, respectively. AGC targets were set to 10^6^ (MS1) and 10^5^ (MS2), normalized collision energy to 28, dynamic exclusion to 10 s, and maximal injection times to 60 ms (MS1) and 120 ms (MS2). Mass spectrometry data were analysed and manually validated using the OpenMS pipeline RNPxl and Open MS TOPPASViewer (27). All crosslinks are listed in **Supplementary Table 3**.

### Surface Plasmon Resonance (SPR)

SPR studies were performed using a BIACORE 200 system (GE Healthcare). tRNA^Gln^ was 3’ biotin labeled with the Pierce ™ RNA 3’ end biotin labeling kit (Thermo Fisher) according to manufacturer’s recommendations. SA-chip (GE-Healthcare) was prepared according to manufacturer’s instruction before biotin labeled RNA diluted in DEPC-H2O was coupled. Analysis of RNA-protein interactions was performed at a flow rate of 30 µL/min in running buffer (150 mM NaCl, 50 mM HEPES-NaOH 7.5, 0.05 % (v/v) Tween) and at 25 °C and 10 Hz data collection rate. Analyte of interest was diluted in running buffer and single-cycle concentration series were injected on the chip with 240 sec contact time per concentration and a final dissociation time of 900 sec. To remove any residual attached protein after the last analyte injection, 2 x 2 minutes regeneration injections with 0.5 % (w/v) SDS were done in between runs. Concentration series reached from 62.5 nM to 4000 nM for hTRMT2A WT and 15.6 – 1000 nM for hTRMT2A CDM (Catalytic death mutant) protein. Data were analyzed with the BIAevaluation software (GE Healthcare). Obtained binding curves were double-referenced against the signal in the empty reference channel and a buffer run. At equilibrium of the binding curves, the corresponding response was plotted against analyte concentration. The KD was determined by fitting this curve to a steady-state affinity model. All experiments were performed in triplicates.

### Proximity-dependent biotinylation

BioID experiments were conducted to identify the hTRMT2A WT and hTRMT2A CDM proximity interactome in HEK 293t cells as described earlier (28) with 5 replicates per condition. At 70 % cell confluency protein expression of BirA*-tagged protein of interest or BirA* tag alone was induced by the addition of doxycycline. 6 h later 1 mL sterile-filtered 20x DMEM/biotin was added to the cells to reach a final concentration of 50 µM biotin. Cells were incubated for 16-18 h. Prior to trypsinization, cells were washed three times with PBS. Detached cells (8x10^7^ to 1x10^8^ per replicate) were centrifuged for 5 min at 4 °C with 300 x *g*. Cells were resuspended in 5 mL lysis buffer (50 mM Tris-HCl pH 7.4, 500 mM NaCl, 0.2 % SDS) freshly supplemented with 1:1000 benzonase (Thermo), 1 mM DTT and Pierce® Protease Inhibitor (Roche). Thereafter, cell suspension was incubated on a rotating wheel at 4 °C for 30 min. After 500 µL of 20 % Triton X-100 was added, the samples were sonicated for 1 min with 30 % duty cycle (Bandelin Sonopuls GM 70UV 70). Three sonication rounds were performed, and the samples were incubated on ice for 5 min in between them. Before the third sonication round, 4.5 mL pre-chilled 50 mM Tris-HCl pH 7.4 was added. Sonicated samples were centrifuged for 10 min at 4 °C with 16,100 x *g*. 500 µL Dynabeads MyOne Streptavidin T7 magnetic beads (Thermo) per condition were equilibrated in 1.5 mL lysis buffer and 1.5 mL 50 mM Tris-HCl, pH 7.4. Beads were placed in a magnetic stand (DYNAL®, Invitrogen). Then, the protein-containing supernatant from the centrifugation step was added to the equilibrated beads. Solution and beads were incubated on a turning wheel overnight at 4 °C. The supernatant was removed from the beads and substituted with 8 mL of wash buffer 1 (2 % SDS in MilliQ-H2O). The wash step with wash buffer 1 was repeated. Next, beads were washed once with wash buffer 2 (0.1 sodium deoxycholate, 1 % Triton X-100, 1 mM EDTA, 500 mM NaCl, 50 mM HEPES-NaOH pH 7.5) and wash buffer 3 (0.5 % sodium deoxycholate, 0.5 % NP-40, 1 mM EDTA, 250 mM LiCl, 10 mM Tris-HCl pH 7.4). To remove detergent, beads were washed three times with wash buffer 4 (1 mM EDTA, 20 mM NaCl, 50 mM Tris-HCl pH 7.4). Washed beads were resuspended in 1.5 mL of 50 mM Tris-HCl pH 7.4 and then put in magnetic stand to remove supernatant. For elution 50 µL 1x sample buffer (50 mM Tris-HCl pH 6.8, 12 % sucrose, 1 % SDS, 0.004 % bromophenol blue, 20 mM DTT) supplemented with 3 mM biotin was added to beads, mixed and incubated at 98 °C for 7 min. Subsequently, samples were placed in magnetic stand, and supernatant subjected to mass spectrometry or analytical Western blot analysis. Enriched and depleted proteins are listed in **Supplementary Table 4**.

### Co-Immunoprecipitation

Co-IP experiments were performed to identify the hTRMT2A proteome in HEK 293t cells with 5 replicates per condition. At 70 % cell confluence protein expression of the protein of interest or FLAG tag alone was induced with doxycycline. 16-18 h later GFP expression was inspected with fluorescence microscopy and cells were washed twice with PBS. For cell harvest, 10 mL ice-cold PBS was added and cells were scraped off the plate. Cell suspension was centrifuged for 5 min at 4 °C with 300 x *g*. Pellets were flash-frozen in liquid nitrogen to ensure equal sample time points.

Cell pellets were thawed and resuspended in 1 mL lysis buffer (50 mM Tris-HCl pH 7.8, 150 mM NaCl, 1 mM EDTA, 1 % Triton X-100) supplemented with one tablet Pierce® Protease Inhibitor (per 100 mL lysis buffer). Suspension was transferred to a new tube and centrifuged for 10 min at 4 °C with 16,100 x *g*. Meanwhile, 50 µL Anti-FLAG® M2 Magnetic Beads (Merck) were equilibrated two times with 900 µL TBS (50 mM Tris-HCl pH 7.8, 150 mM Nacl) and three times with 900 µL lysis buffer. In between wash steps the beads were incubated for 3 min on a magnetic stand (DynaMag 2, DYNAL®, Invitrogen). Then, 3.5 mg protein from the supernatant of the centrifugation step was added to the equilibrated beads. Protein concentration was measured with the Bradford assay solution (Roth) according to instructions. Beads were incubated for 1 h at 4 °C in the shaking mode on a rotator wheel. Thereafter, unbound protein was removed by washing three times with 900 µL high salt wash buffer (50 mM Tris-HCl pH 7.8, 250 mM NaCl, 1 mM EDTA, 1x Triton X-100). The third wash step was incubated for 5 min at 4 °C in shaking mode on a rotator wheel. To remove detergent, beads were washed three times with 900 µL 50 mM Tris-HCl, pH 7.8. Beads were distributed to two tubes for elution: 900 µL for mass spectrometry and 100 µL for Western blot analysis. Elution was performed by addition of 20 µL 2x Laemmli buffer (4 % (w/v) SDS, 20 % glucose, 0.004 % bromophenol blue, 125 mM Tris-HCl pH 6.8) and incubation for 10 min at 98 °C. Finally, tubes were placed into the magnetic stand and the supernatant was transferred to a new tube for respective mass spectrometry analysis and analytical Western blot analysis. Enriched and depleted proteins are listed in **Supplementary Table 5**.

### Western blot analysis

From BioID and Co-IP experiments, the sample was mixed with SDS loading buffer (110 mM Tris-HCl pH 6.8, 40 % (v/v) glycerine, 40 mM DTT, 4 % (w/v) SDS, 0.25 % (w/v) bromophenol blue) and heated for 10 min to 95 °C. Cell lysates from TRMT2A WT and KD cell lines (20-50 µg) were supplemented with Laemmli buffer and boiled for 5 min at 95 °C. Samples were run on SDS-PAGE and transferred onto activated nitrocellulose membrane by semi-dry blotting. The membranes were blocked (5% skimmed milk or 1% Casein in TBS-T for 60 min) and incubated with the primary antibody at 4 °C overnight. Membranes were washed three times for 10 min in TBS-T. In case of anti-Streptavidin-HRP antibody (1:10,000; Thermo, #N100) the blot was washed five times for 5 min and thereafter directly visualized. Other blots were incubated with the secondary horse radish peroxidase (HRP) coupled antibody for 1-2 hours at room temperature. Subsequently, the membranes were washed three times in TBS-T for 10 min, and the chemiluminescence signal was detected using the Super Signal® West Femto Maximum Sensitivity Substrate (Thermo Scientific) or Pierce®ECL Western blotting detection kit according to manufacturer’s instructions. Chemiluminescence was visualized using the Fusion SL 4 device (Vilber Lourmat) or the Alliance UVltec system (Biometra) systems. Primary antibodies used: rabbit anti-TRMT2A (1:1000; Sigma #HPA001077), mouse anti-FLAG (1:2000; Sigma #F3165), rat anti- H3 (1:10,000; Abcam #ab1791), rat anti-FLAG (1:10; in-house, IgG1, monoclonal), rabbit anti-PARP1 (1:1000; Sigma-Aldrich HPA045168), mouse anti-GAPDH (1:500, DSHB-hGAPDH, #2G7). Secondary HRP-coupled antibodies used: goat anti-rabbit (1:10,000; Abcam, #ab6721), mouse anti-rat (1:1000; in-house, polyclonal), sheep anti-mouse (1:10,000; GE Healthcare #NXA931V) or donkey anti-rabbit (1:10,000; GE Healthcare #NA934V).

### LS-MS/MS analysis of BioID and Co-IP samples

Proteins were proteolyzed with LysC and trypsin with filter-aided sample preparation procedure (FASP) as described (29, 30). Acidified eluted peptides were analyzed on a QExactive HF-X mass spectrometer (Thermo Fisher Scientific) online coupled to a UItimate 3000 RSLC nano-HPLC (Dionex). Samples were automatically injected and loaded onto the C18 trap cartridge and after 5 min eluted and separated on the C18 analytical column (Acquity UPLC M-Class HSS T3 Column, 1.8 µm, 75 µm x 250 mm; Waters) by a 95 min non-linear acetonitrile gradient at a flow rate of 250 nl/min. MS spectra were recorded at a resolution of 60000 with an automatic gain control (AGC) target of 3e6 and a maximum injection time of 30 ms from 300 to 1500 m/z. From the MS scan, the 15 most abundant peptide ions were selected for fragmentation via HCD with a normalized collision energy of 28, an isolation window of 1.6 m/z, and a dynamic exclusion of 30 s. MS/MS spectra were recorded at a resolution of 15000 with a AGC target of 1e5 and a maximum injection time of 50 ms. Unassigned charges and charges of +1 and >8 were excluded from the precursor selection.

#### Data processing protocol

Raw spectra were imported into Progenesis QI software (version 4.1). After feature alignment and normalization, spectra were exported as Mascot Generic files and searched against the human UniProt database (20,434 sequences) with Mascot (Matrix Science, version 2.6.2) with the following search parameters: 10 ppm peptide mass tolerance and 20 mmu fragment mass tolerance, one missed cleavage allowed, carbamidomethylation was set as fixed modification, methionine oxidation, and asparagine or glutamine deamidation were allowed as variable modifications. A Mascot-integrated decoy database search calculated an average false discovery of <1% when searches were performed with a mascot percolator score cut-off of 13 and an appropriate significance threshold p. peptide assignment was re-imported into the Progenesis QI software and the abundances of all unique peptides allocated to each protein were summed up. The resulting normalized abundances of the individual proteins were used for the calculation of fold-changes of protein ratios between conditions. Proteins with a sample/control ratio of at least 4 were considered differentially abundant. Statistical analysis was performed on log2 transformed normalized abundance values using Student’s t-test. Changes in protein expression between conditions were considered significant at p < 0.05. Principal component analysis was performed in Progenesis QI software.

### GO-term analysis

„Biological processes“ (BP) and „cellular compartments“ (CC) gene ontology (GO) term analysis was performed with data from BioID experiments using the GO enrichment analysis tool from gene ontology using the PANTHER databank (31). UniProt protein names of significantly enriched hits from BioID experiments were used for GO enrichment analysis. Significance of GO-term enrichment was calculated as false discovery rate (FDR) with Fisher’s exact test. Using custom R-scripts data were visualized as enrichment of respective GO-term on x-axis, the significance of enrichment with color code, and the counts per GO-term with node size.

### Generation of BioID and Co-IP stably transfected cell lines

For stable transfection of HEK 293t cells a piggyBac (PB) transposon plasmid was used (32, 33). Co-transfection with the PB helper plasmid, encoding for PB transposase, allows for stable insertion of the gene of interest into the chromosome. For transfection, HEK 293t cells were plated in a 6-well plate and grown to 80 % confluency. According to the Lipofectamine™ 3000 transfection kit (Invitrogen), 2.5 µg of PB plasmid with the gene of interest, 2.5 µg helper plasmid were transfected into HEK 293t cells and incubated for 3 days. For antibiotic selection complete medium (Gibco^TM^ DMEM + 10 % FBS + 1 % PenStrep) supplemented with hygromycin B (0.7 mg/mL, Invitrogen) was added and exchanged 2-3 times a week for stable integration of the PB plasmid into the genome.

The same expression level of the protein of interest was achieved by fluorescence-activated cell sorting (FACS). Protein expression in cells was induced with doxycycline 18 h before FACS experiment. For FACS the cells were detached with trpysin, washed twice with PBS for 5 min and resuspended in sterile FACS buffer (0.5 % BSA, 2 mM EDTA, 25 mM HEPES-NaOH pH 7.5 in PBS). Directly prior to FACS experiment, the cells were passed through a 35 µm cell strainer. First gating step subtracted dead cells and aggregates. A second gate was set to sort the cell population with a low GFP signal. Sorted cells were plated in 6-well plates and propagated with complete medium containing hygromycin-B.

### Generation of stable RNAi-mediated KD of hTRMT2A

Stable RNAi-mediated knockdown of hTRMT2A was achieved as described earlier (34). HEK 293t cells were infected with commercially available Lentiviral particles for hTRMT2A KD and scrambled control cell line (MISSION® shRNA Lentviral Transduction Particles: scrambled control WT = SHC002V, KD1 = NM_182984.2-**856**s1c1, KD2 = NM_182984.2-**1574**s1c1; Sigma-Aldrich). Cells with stable integration of shRNA were selected as puromycin-resistant colonies. hTRMT2A silencing effect was probed by Western blotting.

### Ribosome isolation and mass spectrometry analysis of rRNA modification status

Ribosome isolation from TRMT2A WT, KD1 and KD2 HEK 293t cells was performed according to a protocol published earlier (35). Briefly, 80 % confluent cells were harvested in ice cold PBS and centrifugated at 4 °C. Cell pellet was resuspended in equivolume buffer A (250 mM sucrose, 250 mM KCl, 5 mM MgCl2, 50 mM Tris-HCl pH 7.4). For cell lysis 0.7 % (v/v) NP-40 was added and incubated on ice for 10-15 min. To separate nuclei, cell lysate was centrifugated for 10 min at 750 x g and 4 °C. To prepare fraction without mitochondria, the supernatant was centrifugated for 10 min at 12,500 x g and 4 °C. Post-mitochondrial fraction (PTM) supernatant was transferred to a new tube and the KCl concentration was adjusted to 500 mM using a 4 M stock solution. Adjusted PTM was carefully layered on top of a 2 mL sucrose cushion (1 M sucrose, 500 mM KCl, 4 mM MgCl2, 50 mM Tris-HCl pH 7.5). Samples were ultracentrifuged for 2 h at 250,000 x g and 4 °C. Supernatant was discarded and translucent ribosome pellet was washed with 200 µL cold DEPC-treated H2O. Finally, ribosomes were resuspended in 3x 100 µL buffer C (25 mM KCl, 5 mM MgCl2, 50 mM Tris-HCl pH 7.4).

For rRNA isolation, 100 µL of ribosomes diluted in buffer C, were mixed with 15 µL DEPC-H2O and 15 µL 3 M NaOAc. Next, 1 volume phenol-chloroform-isoamylacohol (25:24:1) was added. After centrifugation for 1 min at 16,100 x *g*, upper aqueous phase was mixed with 1 volume chloroform and again centrifugated. The recovered upper phase was precipitated in presence of 0.5 µL RNA-grade glycogen and 2 volumes ethanol for > 20 min at -20 °C. RNA was pelleted for > 20 min at 16,100 x *g* and 4 °C. Dry pellet was resuspended in 20 µL DEPC-H2O.

For isolation of 18S and 28S rRNA, purified rRNA was separated on 0.8 % agarose gel for 80 min at 120 V in 1 x TAE buffer. The bands for 18S and 28S rRNA were cut from the gel and electroeluted for 200 min at 150 V in 1 x TAE buffer. After electro-elution the rRNA was precipitated with 2 volumes of 100 % ethanol for 30 min at -20°C and further pelleted for 30 min at 13,200 rpm at 4 °C. To increase the yield, 1 µL of RNA-grade glycogen per 1 mL of the solution was added. The pellet was dissolved in water and the samples were frozen in liquid nitrogen for further storage in -80°C. All reagents used were RNAse and sodium ions free. Integrity and purity of rRNA was checked with TapeStation analysis done using the Agilent TapeStation 4150 with the Agilent High Sensitivity RNA ScreenTapes according to manufacturer’s instructions.

### rRNA digestion to the nucleoside level

500 ng – 5 µg of pre-purified rRNA subunits (20 µL of each) were digested to single nucleosides by using 2 U benzonase nuclease (> 90 % purity, Merck) 0.2 U phosphodiesterase I (VWR), and 2 U alkaline phosphatase (QuickCIP, NEB) in MgCl2 (1 mM) containing Tris (pH 8, 5 mM) buffer. In a final volume of 30 µL 1 µg pentostatin, 5 µg tetrahydrouridine and a final concentration of 10 µM butylated hydroxytoluene were added to each reaction for protection against degradation of the released nucleosides. After incubation for 2 h at 37 °C, 20 µL of LC-MS Buffer A (see below) was added to the mixture and then filtered through 0.2 µm Supor Natural PP filters (Pall Corporation, AcroPrep Advance350, 96 well-plate) at 3000 x g and 4 °C for 30 min before measurement by LC-MS/QQQ.

### LC-MS/QQQ analysis of nucleosides

For quantitative mass spectrometry an Agilent 1290 Infinity equipped with a variable wavelength detector (VWD) combined with an Agilent Technologies G6490 Triple Quad LC/MS system with electrospray ionization (ESI-MS, Agilent Jetstream) was used. Operating parameters: positive-ion mode, cell accelerator voltage of 5 V, N2 gas temperature of 120 °C and N2 gas flow of 11 L/min, sheath gas (N2) temperature of 280 °C with a flow of 11 µL/min, capillary voltage of 3000 V, nozzle voltage of 0 V, nebulizer at 60 psi, high-pressure RF at 100 V and low-pressure RF at 60 V. The instrument was operated in dynamic MRM mode (**Supplementary Table 6**). For separation an Uptisphere C18-HDO column (3.0 µm, 150 x 2.1 mm from Interchim, UP3HDO-150/021) was used. Running conditions were 35 °C and a flow rate of 0.35 mL/min in combination with a binary mobile phase of 5 mM NH4OAc aqueous buffer A, brought to pH 4.9 with glacial acetic acid (200 µL/L), and an organic buffer B of 2 mM NH4COOH in 80 % acetonitrile (Roth, Ultra LC-MS grade, purity ≥99.98). The gradient started at 100 % solvent A for 0.5 min, followed by an increase of solvent B to 10 % over 5.5 min. From 6.0 min to 8.5 min, solvent B was increased to 20 % then to 80 % in 1 min and maintained at 80 % for 1.5 min before returning to 100 % solvent A in 0.5 min and a 2.2 min re-equilibration period. Of each sample 10 µL were co-injected with 1 µL of stable isotope labeled internal standard (ISTD) which was aspirated automatically before each injection from the instrument itself. The sample data were analyzed by the quantitative MassHunter Software from Agilent using the integrated calibration function. The calibration solutions ranged from 0.05 pmol to 100 pmol for each canonical nucleoside and from 0.002 pmol to 5 pmol for each modified nucleoside (12 calibration levels, 1:2 dilution).

The measured molar amount of each canonical nucleoside (pmol) was divided by their expected occurrence in the respective rRNA subunit reported in literature (# per molecule from modomics (4)). 18S rRNA: #C = 493, #U = 356, #G = 540, #A = 403 (RNA_Source_ID = J01866) 28S rRNA: #C = 1646, #U = 687, #G = 1790, #A = 781 (RNA_Source_ID = K03432) Like that the measured amount of injected rRNA subunit in pmol is calculated by each of the four canonical nucleosides separately. Those results were averaged and used as normalization for the modifications to receive the number of modifications per rRNA molecule in the respective sample.

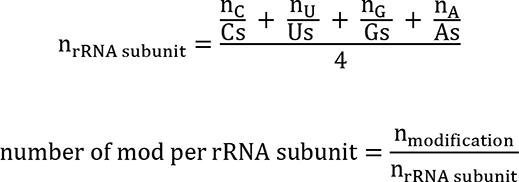

### Immunostainings

Immunocytochemistry (ICC) experiments were performed to determine hTRMT2A localization in HEK 293t cells. Cells on coverslips were fixed for 10 min at RT using 3.7 % formaldehyde solution in PBS. After two PBS washing steps, cells were permeabilized for 5 min with 0.5 % Triton X-100 in PBS. Next, cells were blocked for 30 min using blocking buffer (1 % goat serum in PBS-T). Primary antibody diluted in PBS was added for 1 h at RT. After three washing steps with PBS-T, cells were incubated with secondary antibody diluted in blocking buffer for 1 h at RT in the dark. Cells were washed three times with PBS-T. Next, the cell nuclei were stained with DAPI at a concentration of 0.5 µg/mL DAPI (Thermo) in PBS for 5 min at RT and washed two times with PBS. Finally, coverslips were mounted using the ProLong™ Diamond Antifade Mountant (Invitrogen). Cells were imaged with the Leica DMi8 fluorescence microscope and the Leica imaging software. Images were analysed with the Fiji ImageJ software. Primary antibody used: mouse anti-BirA (1:10; Novus Biologicals, #5B11c3-3). Secondary antibody used: goat anti-Mouse IgG Alexa Fluor 647 (1:2000; Invitrogen, #A-21235)

### Dual luciferase translation assay

The dual Luciferase assay to probe translation fidelity was performed using three Renilla-Firefly fusion protein constructs. In two Renilla sequences mutations were introduced at residues crucial for luciferase activity (36).

hTRMT2A WT (scrambled control) as well as KD1 and KD2 (scrambled control) were cultured (Gibco^TM^ DMEM + 10 % Gibco^TM^ FBS + 1 % Gibco^TM^ Antibiotic-Antimycotic) and seeded at a density of 20,000 cells per well in a 96-well-plate suitable for luminescence measurements (greiner bio-one #655075). Cells were transfected with Renilla and Firefly fusion constructs (WT, D120N, E145Q) using *Trans*IT®- LT1 Transfection Reagent (Mirus Bio) according to the manufactureŕs instructions. After 48 h of incubation at 37 °C and 5 % CO2 culture medium was exchanged with fresh growth medium and Luciferase activity was measured with the Dual-Glo® Luciferase Assay Kit (Promega) according to manufactureŕs instructions using the Tecan Spark® plate reader. Integration time was 10 s. Data represent mean ± SD of 6 biological replicates, in technical triplicates. Statistical analysis was performed with the Wilcoxon rank test and student’s t-test in R.

## Results

### TRMT2A shows low binding specificity for its physiological target tRNAs

To understand how specificity of tRNA methylation is achieved by hTRMT2A, it is imperative to assess both, RNA binding and catalytic activity of hTRMT2A (10). For *in vitro* characterization of the TRMT2A-binding preferences, we prepared recombinant human TRMT2A full-length (hTRMT2A FL, **Supplementary Figure 1B**) and different *in vitro* transcribed RNAs (**Supplementary Figure 1A**). We assessed binding with an Electrophoretic Mobility Shift Assay (EMSA), where ^32^P-labelled RNA was preincubated with increasing concentrations of hTRMT2A FL and polyU as a competitor to preclude non-specific binding. To explore how important the uridine in position 54 of a given tRNA is for binding, we included two tRNAs with U54 (tRNA^Gln^, tRNA^Phe^) and two tRNAs lacking U54. The latter consisted of tRNA^Ala^ with A54 and a mutant version of tRNA^Gln^ where U54 had been changed into G (tRNA^Gln^ ^U54G^). hTRMT2A FL bound all these tRNA with apparent equilibrium dissociation constants (KD) in a low nanomolar range (**Figure 1A**). A somewhat stronger binding was observed for tRNA^Gln^, where a shifted band was already observed at a protein concentration of 60 nM. Of note, for all tRNAs except for tRNA^Phe^ we observed supershifts that imply multiple hTRMT2A enzymes to bind to respective RNA. However, since these supershifts only occurred at higher, micro-molar protein concentrations, we assume that they represent unspecific oligomerization without physiological importance. Also, for the negative control polyU-RNA the binding appeared only as a smear and not as specific band shift, indicative of a rather unspecific interaction (**Figure 1B**).

**Figure 1.**
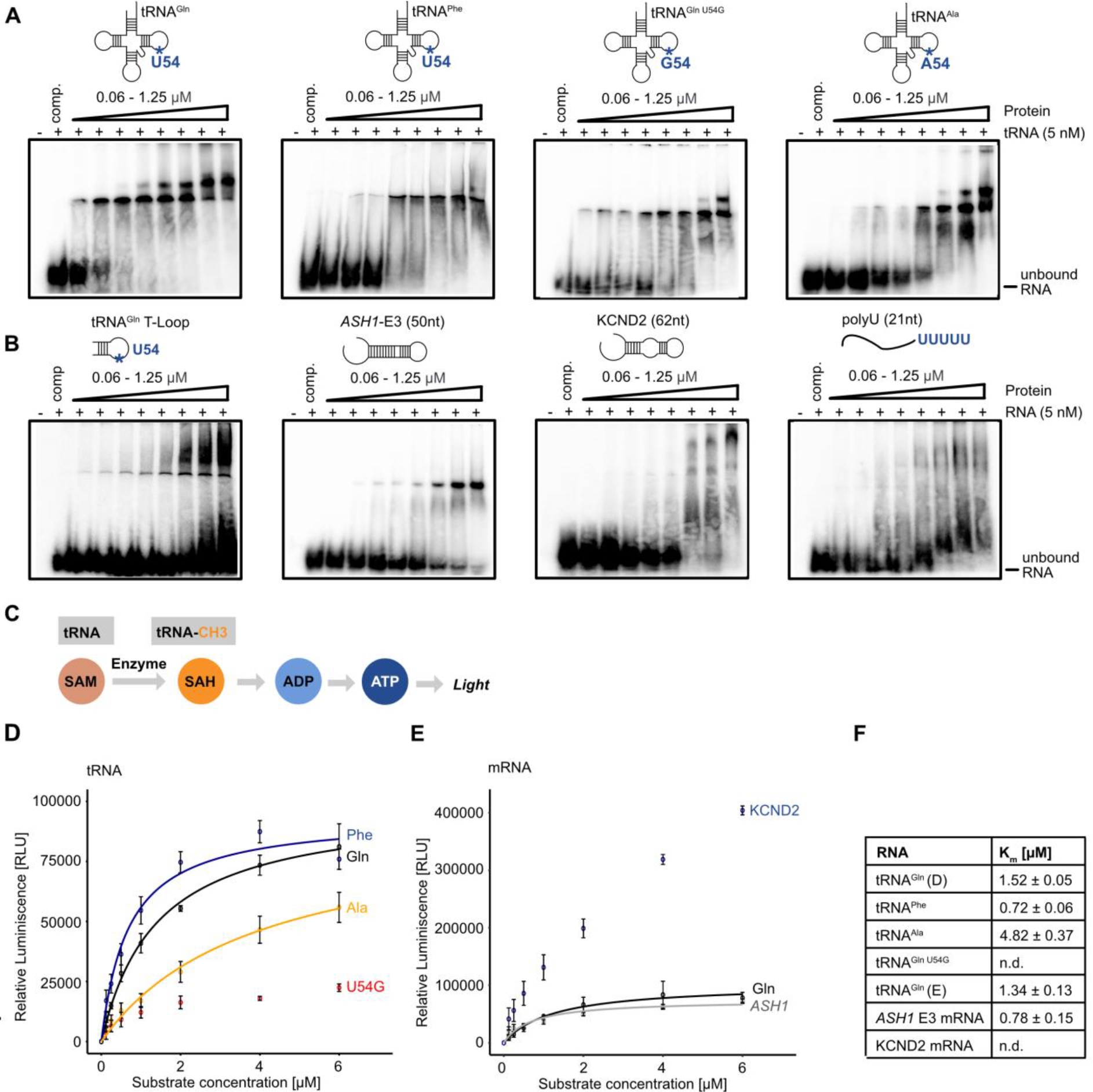
hTRMT2A FL (aa 1-625) binds and methylates RNA with low specificity. **A** EMSAs of tRNA with differing nucleotides at position 54 shows for all tRNA high affinity binding in low nanomolar KD range. **B** EMSA of truncated tRNA^Gln^ T-loop displays unspecific binding with unbound RNA at high protein concentrations. EMSA with mRNA derived from *S. cerevisiae ASH1* E3 element (1771-1821; 51nt) shows binding in nanomolar KD range. EMSA with mRNA derived from human KCND2 (chr7: 366813-366875; 62nt) shows binding in micromolar KD range, whereas control experiment using polyU (21 nt) shows no binding. Binding assays were performed with ^32^P labelled RNA. For experiments polyU competitor and labelled RNA were pre-incubated with increasing concentrations of hTRMT2A FL for 20 min. Reaction without protein served as control. Free RNA and RNA-protein complex were separated on 4 % native PAGE. Each experiment was performed in triplicates. **C** Principle of the MTase Glo™ Methyltransferase assay for methylation experiments. **D** Results of tRNA methylation experiments. **E** Results of mRNA methylation experiments with tRNA^Gln^ as control. Data represent mean ± SD for three replicates. Curves were fitted with Michaelis-Menten equation illustrated as lines. **F** Listed Km values from Michaelis-Menten fitting. For methylation experiments hTRMT2A FL and methyl-donor SAM were preincubated with increasing concentrations of *in vitro* transcribed RNAs for 1 h. Reaction without RNA served as control.

To dissect the structural requirements for hTRMT2A FL binding, we also performed EMSAs with the tRNA^Gln^ T-Loop and two structured fragments derived from mRNA: a fragment of the KCND2 mRNA, which consists of the previously published stem-loop target of hTRMT2A (10). Moreover, the unrelated E3 stem-loop of the *S. cerevisiae ASH1* mRNA, which is not known to be methylated by hTRMT2A, but contains the previously identified sequence motif in the predicted loop recognized by hTRMT2A (**Supplementary Figure 2C**) (10, 37). The tRNA^Gln^ T- Loop showed a weaker shift with most of the RNA remaining in its free form, which suggests the formation of complexes with lower stability (**Figure 1B**). Both stem-loop containing mRNAs displayed binding, however in a high nanomolar to micromolar KD range (**Figure 1B**), which indicates lower affinity than observed with tRNAs. KCND2 showed a somewhat less defined binding than *ASH1* E3, indicated by the lack of a distinctly shifted band. However, an additional band at higher molecular weight could point into the direction of a secondary binding event.

In summary, these results suggest that hTRMT2A FL binding to tRNAs is independent of the presence of a U54 as a potential methylation site and that the enzyme binds *in vitro* stem-loop structured RNAs with modestly lower affinities than tRNAs. While binding affinities are in the expected range (15), binding specificity of hTRMT2A for its physiological methylation targets appears to be low *in vitro*.

### Enzymatic activity of hTRMT2A FL

To correlate our binding studies with enzymatic activities, we tested the methyltransferase activity of hTRMT2A FL on selected RNAs using the luminescence-based MTase Glo™ assay (**Figure 1C**). In this assay, relative luminescence correlates directly with enzymatic activity. hTRMT2A FL was incubated with increasing RNA concentrations to generate saturated Km curves for Michaelis-Menten fitting. A reaction without RNA was used as background control. We observed that hTRMT2A methylates tRNA^Gln^ and tRNA^Phe^ with a Km of 1.52 ± 0.05 µM and 0.72 ± 0.06 µM, respectively, which is in the range of previously reported Km values for methyltransferases (38). In contrast, the two tRNAs lacking U54 (tRNA^Ala^, tRNA^Gln^ ^U54G^) were methylated orders of magnitude less efficiently (**Figure 1D**). Taken together, binding assays and enzymatic assays indicate that the specificity of target-tRNA methylation is mainly mediated by a uridine residue in the right position and not by the affinity to these tRNAs per se.

KCND2 mRNA experiments resulted in higher overall luminescence, potentially deriving from a secondary methylation site *in vitro* (**Supplementary Figure 2B**). In agreement with unspecific binding (**Figure 1B**), however, KCDN2 mRNA was required in high concentrations and did not reach saturation indicating non-physiologic enzymatic activity. The presence of the RNA-recognition motif of hTRMT2A in the unrelated, stem-loop structured *ASH1* E3 mRNA was sufficient for similar methylation like those of tRNA substrates with a Km of 0.78 ± 0.15 (**Figure 1D, Supplementary Figure 2**). Thus, hTRMT2A cannot only methylate substrate tRNA, but also other stem-loop structured RNAs.

### tRNA is bound by hTRMT2A via multidomain interactions

Based on the results from binding and methylation experiments we choose tRNA^Gln^ to dissect the RNA-interacting domains of hTRMT2A. For this purpose, we generated several hTRMT2A fragments, whose domain borders were chosen based on experimental evidence (34), sequence alignments, and homology models. We purified hTRMT2A FL, the RNA-binding domain (RBD, aa 69-147), and the Central domain (Central, aa 237-414). Unfortunately, the predicted entire methyltransferase domain (aa 179-581) was unsuitable for *in vitro* studies due to protein degradation. Instead, we created a more stable protein fragment, in which the RBD is fused directly to a part of the methyltransferase domain (Fusion, 69-240/413-590) (**Figure 2A**, **Supplementary Figure 1C/D**). In EMSAs, all fragments showed only weak mobility shifts in the micromolar protein range (**Figure 2C**) and hence much weaker binding than hTRMT2A FL (**Figure 2B**). This indicates that none of the individual domains - RBD, Central, and Fusion – are sufficient for full tRNA^Gln^ binding and that they might have to interact cooperatively for physiological binding affinity.

**Figure 2.**
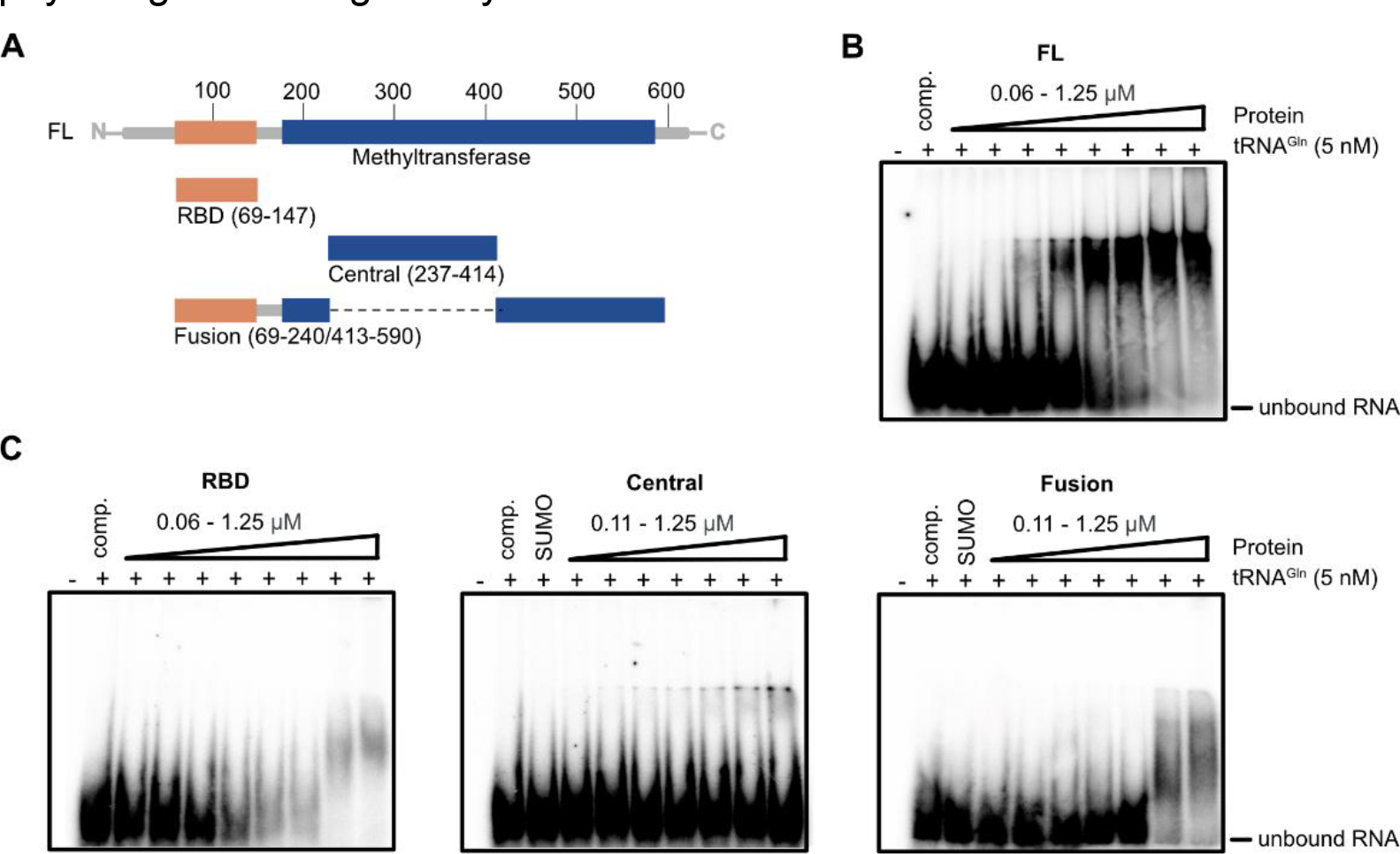
hTRMT2A domains bind tRNA^Gln^ cooperatively. A. Cartoon depicting protein fragments of hTRMT2A used for EMSAs. **B** EMSA with hTRMT2A FL shows high affinity binding to tRNA^Gln^ in nanomolar KD range. **C** EMSAs with hTRMT2A RBD, Central and Fusion domain display only low affinity binding to tRNA^Gln^ in micromolar KD range. Because the SUMO-tagged Central domain was used, a SUMO-tag control was included. Binding assays were performed with ^32^P labelled tRNA^Gln^. For experiments polyU competitor and labelled RNA were pre-incubated with increasing concentrations of hTRMT2A proteins for 20 min. Reaction without protein served as control. Free RNA and RNA-protein complex were separated on 4 % native PAGE. Each experiment was performed in triplicates.

### tRNA binding requires interactions with all hTRMT2A domains

To map the hTRMT2A-tRNA^Gln^ interaction surface with near-amino acid resolution, we performed nitrogen mustard (NM) and ultraviolet radiation (UV) induced cross-linking of the hTRMT2A-tRNA^Gln^ complex followed by mass spectrometry. Experiments were conducted with the *in vitro* reconstituted protein-RNA complex (**Supplementary Figure 3A**). To confirm that hTRMT2A FL forms a stable complex with tRNA^Gln^, we also performed size exclusion chromatography with tRNA^Gln^ and hTRMT2A FL separately and with the hTRMT2A-tRNA^Gln^ complex (1:1 ratio). Stable complex formation was apparent as a shift of the elution peak to smaller elution volumes (from 17.41 mL to 15.46 mL), corresponding to a higher molecular weight (**Figure 3A**). These co-purified complexes were also assessed by mass-spectrometric approaches (**Figure 3A**, **Supplementary Figure 3B**).

**Figure 3.**
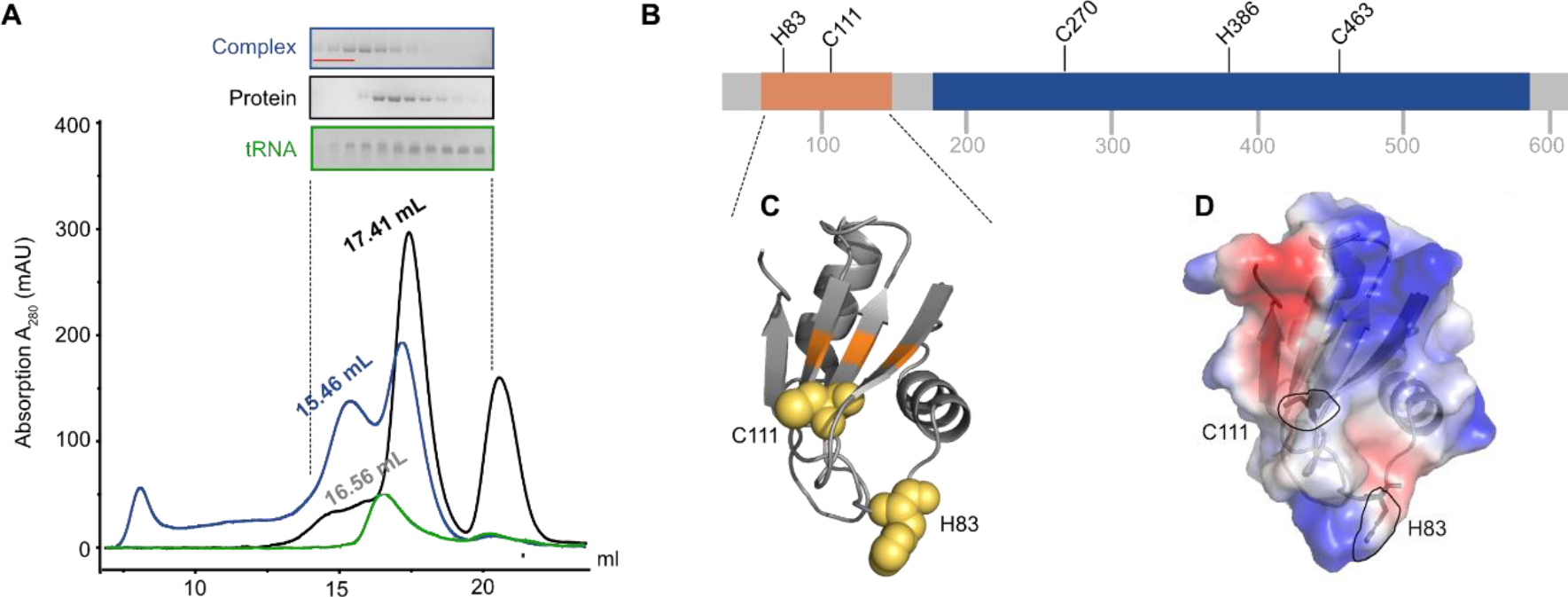
tRNA^Gln^ contacts various amino acids from all hTRMT2A domains. A. Size exclusion chromatography profile from a size-exclusion chromatography run with an S6 (15/150) column with hTRMT2A FL and tRNA^Gln^, as well as the stable protein-RNA 1:1 complex. SDS-PAGE and denaturing PAGE of protein, RNA and complex are shown. Pooled fractions for crosslinking experiments are highlighted in red. **B** Crosslinked amino acids are mapped on a linear representation of hTRMT2A FL and those consistently crosslinked in three or four of four datasets are shown. **C** Crosslinked amino acids (yellow) are mapped on the hTRMT2A RBD X-ray structure (PDB ID: 7NTO) and the predicted RNA binding surface is colored orange. **D** Electrostatic surface potential of hTRMT2A RBD and the position of crosslinked amino acids H83 and C111 (dashed lines, stick representation). Positive (blue) and negative (red) charge profile was prepared with the PyMOL APBS plugin. Figures were prepared with PyMOL (version 2.0.4).

The *in vitro* reconstituted and co-purified samples were cross-linked, trypsin- and RNase-digested, and analyzed by mass spectrometry. Out of the four collected datasets (UV, NM, *in vitro* reconstituted, purified complex) we consistently found in at least three of them the following amino acids to be cross-linked to RNA (**Supplementary Table 3**): C270 to a C base, C463 to A or U, H386 to U/C or G. Both C111 and H83 cross-linked to all four nucleotides (**Figure 3B**). In some instances, these amino acids were also crosslinked to incompletely digested dinucleotides such as UU. Mapping of both crosslinked amino acids C111 and H83 onto the previously published X-ray structure of the RBD from hTRMT2A FL ((34) PDB-ID: 7NTO) shows tRNA interaction with the RBD ß-sheet surface (C111) and with one of the flexible loops (H83) (**Figure 3C**). For RBDs, the ß-sheet surface has been identified as the major RNA binding platform, although also loop-mediated RBD-RNA interactions have been reported (39, 40). The electrostatic surface potential illustrates the position of crosslinked C111 and H83 in relation to the surface charge, where especially the aromatic ring of H83 is part of a positively charged patch (**Figure 3D**).

The cross-linked amino acids C270, H386 and C463 are distributed across the methyltransferase domain. A sequence alignment with *E. coli* TrmA (**Supplementary Figure 4**) revealed that C463 of hTRMT2A corresponds to A241 of *E. coli* TrmA. In the *E. coli* TrmA T-Loop co-structure ((19), PDB ID: 3BT7) residue A241 is close to the catalytic center and in proximity (below 10 Angstrom) to the stem of the T-Loop. Proximal to the crosslinked residue C270 lies G263, which corresponds to R51 of *E. coli* TrmA, a residue shown to modestly contribute to tRNA binding (15).

In two of four datasets 17 additional cross-linked residues were identified (**Supplementary Figure 3C**). Most notably, amongst them is the catalytic cysteine C538 cross-linked to uridine. Since C538 is known to make a covalent bond to the U54 base during catalysis (10, 19), this highlights the specificity of our crosslinking mass spectrometry data. Taken together, these results cross-validate our previous observation from EMSAs that multiple domains are involved in and required for RNA binding.

### Catalytic death mutant of hTRMT2A FL

To better dissect the function of hTRMT2A, we decided to create a mutant version of this enzyme with impaired enzymatic activity. Based on the EMSAs with subdomains of hTRMT2A (**Figure 2**) and our cross-linking experiments (**Figure 3**), we anticipated that an enzymatic activity mutant of hTRMT2A would require surface mutations on the RBD to reduce interaction with target RNA and across the methyltransferase domain to abolish catalytic activity. Thus, these areas were chosen for mutation. For selecting the exact amino acids, sequence conservation (**Supplementary Figure 4**) and the reported impact of mutations on RNA binding in *E. coli* TrmA were considered (18). Finally, we selected three, conserved and solvent-exposed amino acids on the ß-sheet surface of the RBD (E76A, K104E, F113A) and five in the methyltransferase domain including the main catalytic base C538 for mutation (K242A, F409A, Q411A, D510A, C538A). This hTRMT2A mutant version was termed <u>c</u>atalytic <u>d</u>eath <u>m</u>utant (CDM). hTRMT2A CDM was used in comparative RNA-binding and enzymatic activity experiments with hTRMT2A WT, the previously reported catalytic base mutant E581A (CBM), and the catalytic nucleophile mutant C538A (CNM) (10) (**Supplementary Figure 1B**). In EMSAs, CBM and CNM mutants showed slightly less binding to tRNA^Gln^ than WT protein when comparing their fractions of unbound RNA at higher protein concentrations (**Figure 4A**). Surprisingly, the CDM mutant displayed similar binding as WT, but an increase in super-shifted binding events. This highlights the difficulty to abolish RNA binding of a protein with a large interaction surface such as hTRMT2A, reported through our cross-linking experiments.

**Figure 4.**
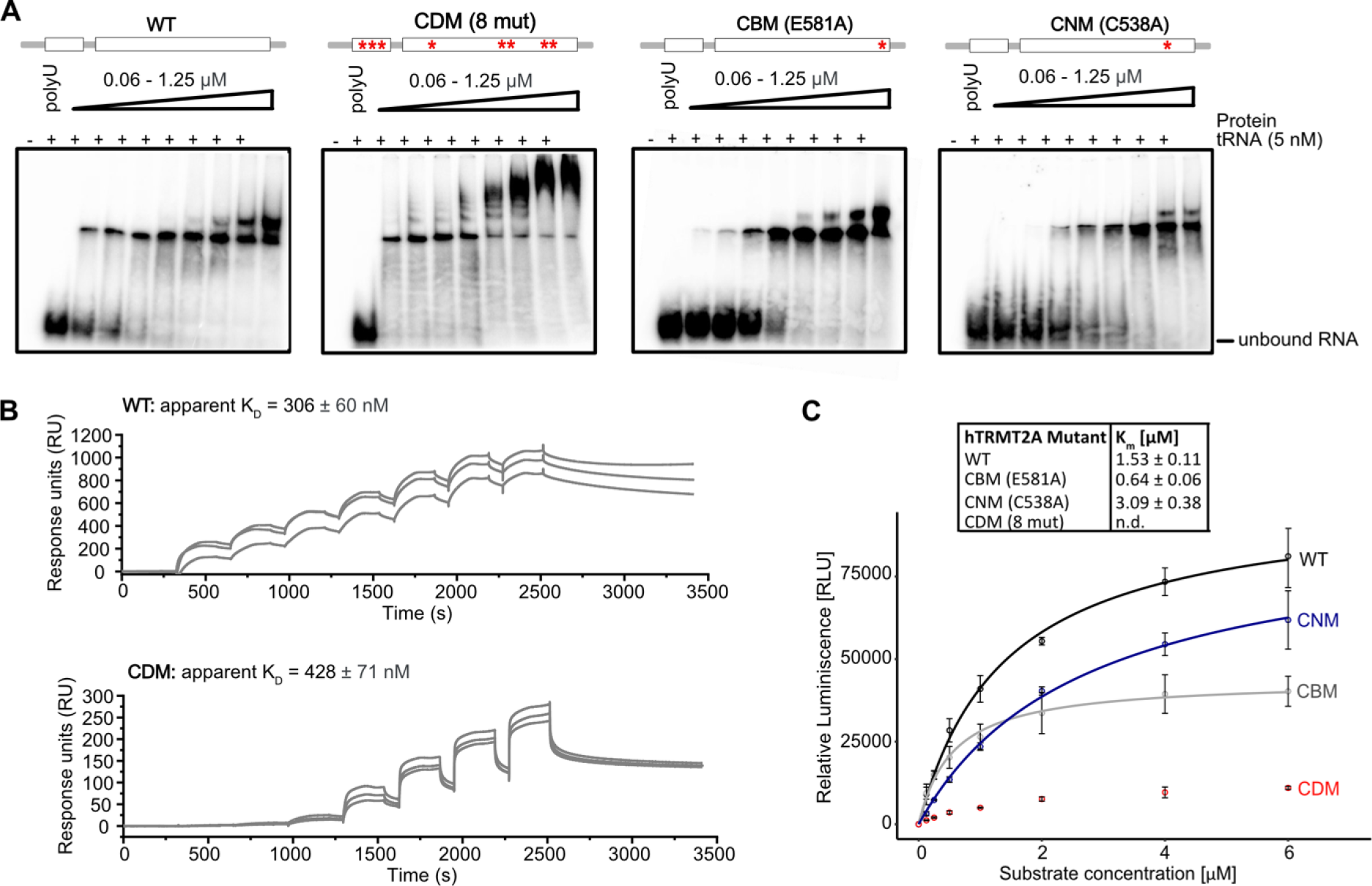
hTRMT2A CDM mutant exhibits impaired enzymatic activity. A. Comparative EMSAs with hTRMT2A WT, CDM, CBM and CNM display similar high affinity binding profiles to tRNA^Gln^ in nanomolar KD range. hTRMT2A CDM shows supershifted binding events. Binding assays were performed with ^32^P labelled tRNA^Gln^. For experiments, polyU competitor and labelled RNA were pre-incubated with increasing concentrations of hTRMT2A for 20 min. Reaction without protein served as control. Free RNA and RNA- protein complex were separated on 4% native PAGE. Each experiment was performed in triplicates. **B** Single-cycle SPR results show similar KD for hTRMT2A WT and hTRMT2A CDM. For all experiments biotinylated tRNA^Gln^ was coupled to an SA chip. A protein concentration series from 62.4 to 4000 nM (WT) and 15.6 to 1000 nM (CDM) was injected resulting in the illustrated, double-referenced curves. Triplicate measurements are shown and respective apparent KD values are shown. More details are available in **Supplementary** Figure 5. **C** Km curve results for hTRMT2A WT, CDM, CBM, CNM mutants from methylation assays confirm strongly reduced enzymatic activity of hTRMT2A CDM mutant. For experiments constant protein concentrations and increasing tRNA^Gln^ concentrations were used. Data represent mean ± SD of three replicates. Curves were fitted with Michaelis-Menten equation illustrated as lines. Km values are listed.

Since EMSAs only provide an estimate of binding parameters, we quantified binding affinities of hTRMT2A WT and CDM by surface plasmon resonance (SPR) experiments with surface-coupled tRNA^Gln^. The experiments indicated rather complex binding events possibly involving multiple interaction sites, which prevented precise determination of binding kinetics. However, in agreement with EMSAs, hTRMT2A WT exhibited high affinity binding with an apparent KD of 306 ± 60 nM and hTRMT2A CDM showed a similar, albeit slightly lower affinity (apparent KD of 428 ± 71 nM) (**Figure 4B, Supplementary Figure 5**). These results indicate that hTRMT2A CDM exhibits a similar binding affinity to tRNA^Gln^ as compared to hTRMT2A WT. However, despite similar tRNA binding of hTRMT2A CDM, its enzymatic activity is almost abolished as clearly visible from methylation assay results (**Figure 4C**).

Further, methylation by hTRMT2A CBM saturated at lower RNA concentrations and a modestly lower KM of 0.64 ± 0.06 µM when compared to hTRMT2A WT (**Figure 4C**), which is in line with irreversible trapping of RNA and blocking of the catalytic center. The hTRMT2A CNM showed a two times higher Km, which indicates less efficient methylation. Since hTRMT2A CDM showed the strongest reduction of tRNA^Gln^ methylation, but similar RNA binding profiles, we selected this mutant for cellular studies. Moreover, this mutant was of particular interest as the *E. coli* homolog of hTRMT2A, TrmA, was shown to exhibit an enzymatic-activity independent secondary function as tRNA chaperone (15). By including a mutant with impaired catalytic activity but similar tRNA binding affinity in cellular studies we hoped to gain insight into this prospective function for hTRMT2A.

### Proximity ligation based interactome of hTRMT2A

To better understand the cellular functions of hTRMT2A, we aimed at an unbiased assessment of its protein interactome and hence its cellular functions by employing proximity-dependent biotinylation (BioID) (28). In this assay, a protein of interest is fused to the promiscuous biotin ligase BirA* which biotinylates nearby proteins, allowing for the detection of proximal proteins with a distance of up to 10 nm (**Supplementary Figure 6A**). In our study, expression of BirA*-tagged hTRMT2A protein from an overexpression plasmid (**Supplementary Figure 7A**) was induced in biotin-supplemented cells. Cells were lysed, biotinylated proteins were captured using streptavidin-beads and analyzed by mass spectrometry (**Supplementary Figure 8A**). To assess the correct sub-cellular localization of the BirA* fusion proteins to nucleus and cytoplasm (23, 41) (proteinatlas.org), we performed immunostainings against FLAG-tag from the overexpression fusion protein. Immunostainings illustrated that FLAG-tagged hTRMT2A primarily localizes to the nucleus, but also in the cytoplasm. These staining confirm indistinguishable subcellular localization of FLAG-BirA*- tagged, overexpressed hTRMT2A variants (**Supplementary Figure 9**).

BirA* biotinylated many cellular proteins as detected by streptavidin Western blotting (**Supplementary Figure 6B**). The fold change ratios of the mass spectrometry results from WT/BirA* and CDM/BirA* were plotted against significance values (**Figure 5A**). After applying cut-off thresholds (Log2 ratio > 2, -Log10 p-value > 3), 222 and 177 proteins were enriched in WT and CDM samples over BirA* background control, respectively. Both datasets shared a subset of 115 proteins. Since hTRMT2A WT and CDM displayed similar RNA binding, a large overlap of 115 proteins was expected and constitute candidates for interaction partners. GO-term analyses for biological processes and cellular compartments revealed enrichment of rRNA processing and ribosome biogenesis for protein function and suggested a primary localization of the identified proteins in the nucleolus (**Figure 5B**). These GO terms are consistent with the immunostainings of hTRMT2A (**Supplementary Figure 9**) further validating the BioID experiment.

**Figure 5.**
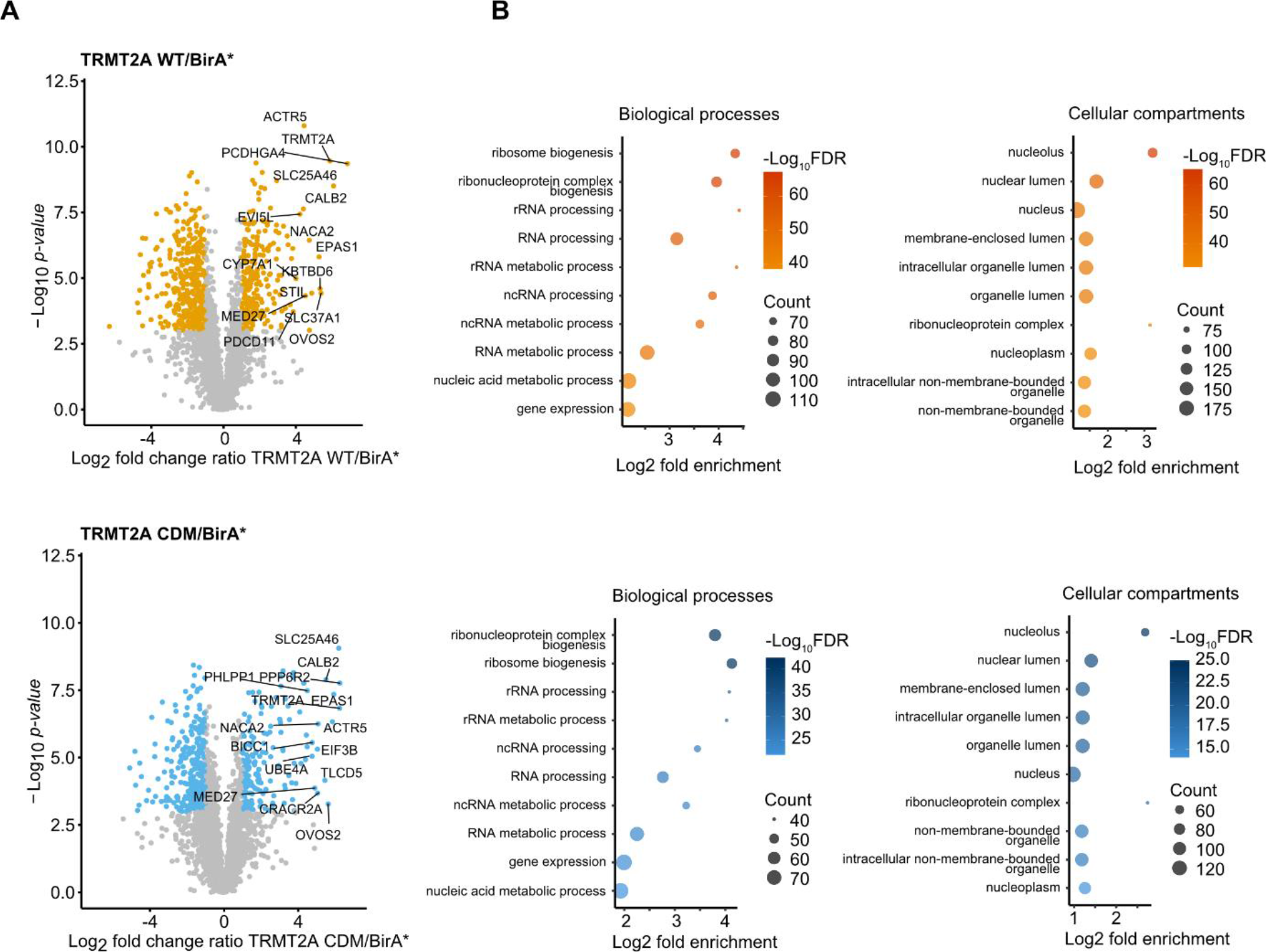
hTRMT2A WT and CDM interactome from BioID experiments. Volcano plots of hTRMT2A WT (**A**) and hTRMT2A CDM BioID (**B**) experiment show enriched proteins and was performed in quintuplicates. Log2 fold change ratio of hTRMT2A WT (BirA*-tagged)/BirA* background and hTRMT2A CDM (BirA*-tagged)/BirA* background was plotted against -Log10 p-value. Ratio cut-offs were > 2 and < 0.5 and significance cut-off, with a p-value < 0.05. Hits in agreement with these thresholds are highlighted yellow (WT) and blue (CDM), respectively. Top 10 hits are labeled with protein names. Top 10 most enriched GO-terms for biological processes and cellular compartments of hTRMT2A WT (**A**, right panel) and hTRMT2A CDM (**B**, right panel) BioID experiments show enrichment of rRNA processing and ribosome biogenesis processes and the nucleolus as most enriched cellular compartment. GO-terms were ranked according to false discovery rate (FDR) as computed with the fisher’s exact test. Graphs illustrate GO-terms plotted against Log2 fold enrichment. Node color represents -Log10 FDR. Node size represents counts per GO-term. GO-terms were visualized in R.

Moreover, we compared differences in the interactome of hTRMT2A WT and CDM by plotting the hTRMT2A CDM/WT ratios against significance levels (**Supplementary Figure 10A**). Notably, proteins that are associated with RNA and nucleic acid metabolic processes were enriched in hTRMT2A WT over CDM. This demonstrates that hTRMT2A CDM interactors are depleted in nucleic acid associated functions, which might be due to its abolished enzymatic inactivity and thus potentially points towards proteins associated with a non- enzymatic, secondary function of hTRMT2A. However, these differences were not very pronounced. Futures studies will be required to dissect if hTRMT2a indeed has such a moonlighting RNA-chaperone function.

### Cross-validation of protein hTRMT2A interactors of by co-immunoprecipitation

To cross-validate the interaction partners identified by BioID with a second proteome-wide approach, co-immunoprecipitation experiments (Co-IP) were performed. Co-IP detects protein interactions that are either direct or protein-/RNA-mediated indirect (**Supplementary Figure 11A**). Expression of FLAG-tagged hTRMT2A protein and an only FLAG-peptide was induced. Cells were lysed, tagged proteins were directly captured using FLAG antibody-coated beads and analyzed by mass-spectrometry (**Supplementary Figure 8B**). Successful elution of FLAG-hTRMT2A and its interacting proteins from beads was confirmed with Western blotting (**Supplementary Figure 11B**). After analysis of the mass spectrometry results, we plotted WT/FLAG and CDM/FLAG fold change ratios against significance values (Log2 ratio > 2, - Log10 p-value > 3). This revealed enrichment of 74 and 72 proteins in hTRMT2A WT and CDM samples over FLAG background control, respectively (**Figure 6A**). To identify potential hTRMT2A interaction candidates, the ratios of significantly enriched proteins from the Co-IP hTRMT2A WT and CDM datasets were plotted against each other. Both datasets shared a subset of 39 proteins (**Supplementary Figure 10B**).

**Figure 6.**
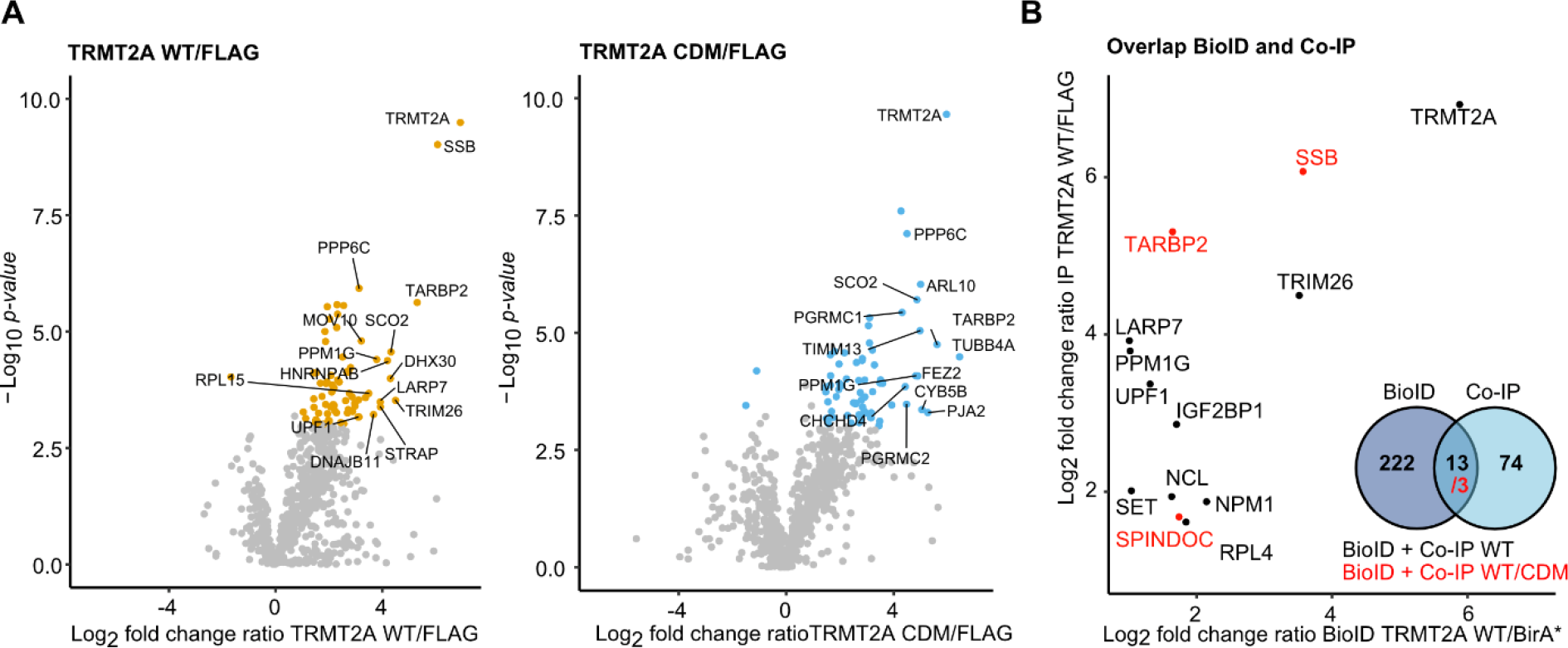
Potential interaction partners of hTRMT2A. A. Volcano plots of hTRMT2A WT and hTRMT2A CDM Co-IP experiment show enriched proteins and was performed in quintuplicates. Log2 fold change ratio of hTRMT2A WT (FLAG-tagged)/FLAG background and hTRMT2A CDM (FLAG-tagged)/FLAG background was plotted against -Log10 p-value. Ratio cut-offs were > 2 and < 0.5 and significance cut-off, with a p-value < 0.05. Hits in agreement with these thresholds are highlighted yellow (WT) and blue (CDM), respectively. Top 10 hits are labeled with protein names. **B** Overlap of significant hits from hTRMT2A WT BioID and Co-IP data shown as plot of the respective ratios (black). Overlap of significant hits from hTRMT2A WT and CDM, BioID and IP experiments (red) shown in the same plot. Venn diagram summarizes overlap of BioID and IP datasets.

The further overlap of hTRMT2A WT data from BioID and Co-IP experiments resulted in 13 common proteins, that constitute confident protein interaction partners (**Figure 6B**). Three of these proteins, SSB (Lupus La protein), TARBP2, and SPINDOC, are also enriched in hTRMT2A CDM data from BioID and Co-IP experiments. These three common proteins constitute cross-validated, highest-confidence interaction candidates of hTRMT2A.

### hTRMT2A does not methylate rRNA

The GO-term annotations for biological processes (BioID) and the identification of ribosomal proteins such as RPL4 or NCL as possible hTRMT2A interactors motivated us to explore a potential functional link between hTRMT2A-dependent methylation and rRNA processing/ribosome biogenesis. Previous cryoEM studies of the human ribosomes had revealed all chemical modifications in rRNAs (42). Two reported sites of m^5^U modifications gained our particular interest: m^5^U4083 in the 60S subunit and m^5^U815 in the 40S subunit. To our knowledge, none of these m^5^U modifications had been previously associated with a dedicated methyltransferase. Recently, FICC-CLIP experiments confirmed a crosslink of hTRMT2A with the small subunit rRNA (10). Moreover, the hTRMT2A homolog hTRMT2B has been shown to methylate tRNA and 12S rRNA in human mitochondria (12). Of note, another mass-spectrometry-based study did not identify m^5^U at respective rRNA positions, wherefore its presence is still discussed (43). Thus, we wanted to clarify whether m^5^U is found at respective cytosolic rRNA positions and whether hTRMT2A is the dedicated m^5^U methyltransferase. For this purpose, two hTRMT2A knockdown HEK 293t cell lines (KD1, KD2) were generated using stable transfection of shRNA. Knockdowns were validated with Western blotting and show a clear reduction of TRMT2A protein levels, with a higher reduction in KD1 (34) (**Supplementary Figure 6B/C**). For experimental assessment of rRNA methylation status, cytosolic ribosomes from hTRMT2A WT, KD1, and KD2 cells were isolated. After rRNA extraction from ribosomes, 18S and 28S rRNA were separated via gel electrophoresis. Tape station analysis showed high RNA purity and effective separation of 18S and 28S rRNA as well as no contamination by tRNA (**Supplementary Figure 12**). Pure 18S and 28S rRNA was subjected to mass-spectrometric analysis and the number of modifications per rRNA was calculated (for details see methods section). Amount of Cm, Um and Ψ modifications per 18S and 28S rRNA correspond well with previously reported numbers (4) (**Supplementary Figure 13A**). Sub-stochiometric m^5^U of 18S and 28S rRNA from hTRMT2A WT was detected in line with around 1 – 4 % methylation of one uridine (**Figure 7B**). This constitutes a much lower level than is usually found for modifications that occur on one site in rRNA (**Figure 7A, Supplementary Figure 13B**). Further, the very low level of m^5^U modifications resembled that of modifications reported to be not present in rRNA (**Supplementary Figure 13C**). Finally, both knockdowns do not further reduce the already low levels of m^5^U modification. In summary, these findings suggest that neither in 18S nor 28S rRNA of HEK 293t cells m^5^U can be found at physiologically relevant levels.

**Figure 7.**
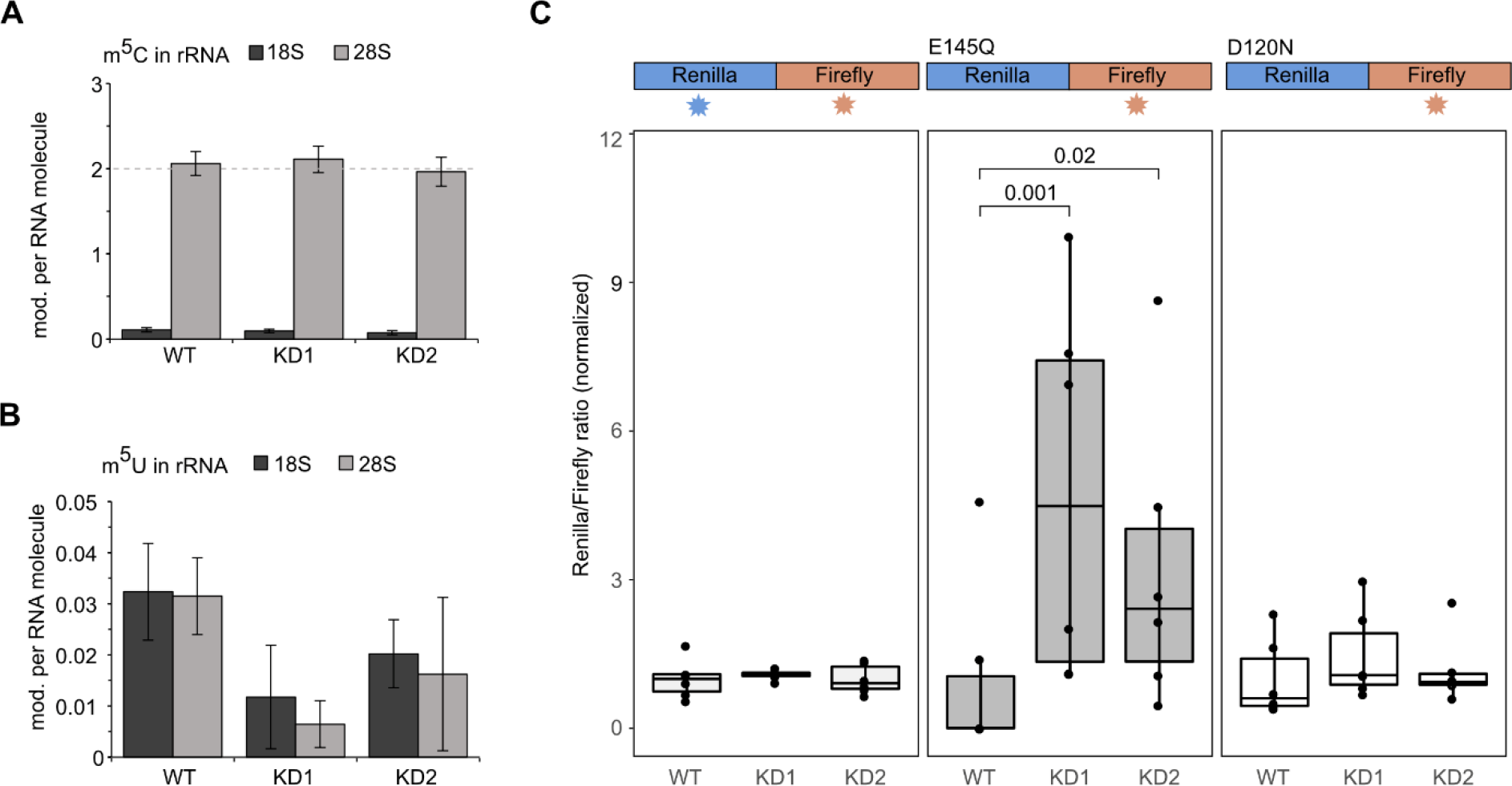
hTRMT2A has not impact on rRNA methylation, but affects translation fidelity A. Barplot of rRNA mass spectrometry analysis of m^5^C in 28S and 18S rRNA shows two modifications in 28S, as reported earlier. **B** Barplot of rRNA mass spectrometry analysis of m^5^U in 28S and 18S rRNA shows very low modification levels that do not change upon hTRMT2A KD. rRNA was obtained from ribosomes that were isolated from hTRMT2A WT and KD1 and KD2 cells. Data represents mean ± SD of three biological replicates. **C** Translation fidelity assay with dual-luciferase WT, E145Q and D120N mutants and their respective luminescence read-out indicated over the graph. Luciferase plasmids were transfected into hTRMT2A WT, KD1 and KD2. For E145Q luciferase mutant, the ratio in KD1 and KD2 was significantly increased over WT, indicating higher translation error rate. WT and D120N luciferase mutant did not show significant changes. Data represents mean ± SD of six biological replicates in technical triplicates. Significance was tested with Wilcoxon test for non-normally distributed values with p > 0.05 = ns, p < 0.05 = significant, p-values are indicated.

### hTRMT2A is important for translation fidelity

As hTRMT2A is the dedicated methyltransferase for tRNA m^5^U formation, lack of this modification was hypothesized to influence tRNA folding and therefore translation (44).

However, for the hTRMT2A paralog hTRMT2B, a lack of m^5^U in tRNA/rRNA did not affect translation efficiency or mito-ribosome stability (12). Indeed, we have recently shown that in a hTRMT2A-deficient situation, protein synthesis is not impaired (34), which suggests normal translation efficiency. Therefore, we asked whether another parameter of translation, translation fidelity, might be affected by hTRMT2A KD. Of note, hypomodification of RNA other than tRNA, such as tRNA fragments (tRF) or mRNA, could impact translation fidelity as well (20, 24, 45).

To test this hypothesis, we used a gain of function reporter previously described to detect small increases in mistranslation rates (36). The reporter construct contains a fusion protein of renilla and firefly luciferases (derived from *Renilla reniformis* and *Photinus pyralis*, respectively). Single amino acid substitutions within the renilla luciferase (E145Q or D120N) render the enzyme inactive. In a condition with increased translation error rate, the renilla luciferase would stochastically be synthesized as a functional protein. The dual luciferase setup allowed us to control for protein expression and therefore measure mistranslation accurately.

The reporter constructs were transfected into hTRMT2A WT and KD cells, and luminescence was measured. Indeed, for the E145Q renilla mutation, both KD1 and KD2 cell lines showed significantly increased normalized luminescence ratios, which are in line with an increased mistranslation in hTRMT2A KD over WT. Of note, a higher increase in luminescence ratios was seen for KD1 cell line, which displayed a stronger reduction of hTRMT2A levels than KD2 in western blot analysis (**Supplementary Figure 7**). However, no significant increase in mistranslation was observed in the same experiment using the D120N mutation (**Figure 7C**). These observations suggest that E145Q is more prone to mistranslation than D120N.

## Discussion

In this study, we characterized the properties of the human tRNA methyltransferase TRMT2A with respect to RNA binding and catalysis. We also identified its RNA-binding interface. Furthermore, we provide the first unbiased assessment of the protein-interaction network of hTRMT2A. These insights suggest novel implications of hTRMT2A and m^5^U modification in translation.

### hTRMT2A is a promiscuous enzyme that binds target RNAs with low specificity

Previously, it has been shown with FICC-CLIP experiments that hTRMT2A binds to the sequence motif GTTCG(A)A (10), which suggests a clear sequence preference of hTRMT2A *in vivo*. In the same study, no further dissection of structural requirements, orthogonal assays, or mutational studies of the sequence motif were performed. Thus, we chose positive hits from these experiments to complement available data with our orthogonal *in vitro* experiments. As an initial result, we observed that the binding affinity of hTRMT2A for tRNAs was in the expected range. The major parameter that increased methylation efficiency was the presence of a uridine positioned in the T-Loop of the tRNA. However, in our experimental setup binding and methylation was not restricted to tRNA as exemplified by the unrelated *ASH1* E3 stem-loop structured RNA containing a hTRMT2A binding motif (**Figure 1**). In conclusion, our *in vitro* data point towards an enzyme, which binds and methylates not only tRNA, but also other structured RNAs, thus displaying no exclusive preference for tRNA.

Previously, hTRMT2A was analyzed regarding its binding (iCLIP) and methylation (FICC- CLIP) of RNA (10). Here, it was apparent that hTRMT2A did not methylate all bound RNA substrates, as exemplified in our data for tRNA^Gln^ ^U54G^ and tRNA^Ala^ binding and methylation (**Figure 1**). These observations could point in the direction of other, m^5^U formation-independent functions of hTRMT2A, for example as a tRNA chaperone (15, 46).

Apart from the consistent results presented here, we reported unspecific binding and methylation of a KCND2 mRNA fragment (10). Hence, it might be that greater specificity is achieved *in vivo* by masking of unspecific methylation sites by other factors or an increase of specificity in RNA binding by co-factors of TRMT2A. Further experiments will be required to clarify whether KCND2 binding, and methylation is indeed a physiologically relevant event *in vivo*.

### *In silico* model of hTRMT2A-tRNA^Gln^ complex supported by experimental evidence

So far, there is no complete structure model of hTRMT2As available. Our data from crosslinking experiments followed by mass-spectrometry resolved the hTRMT2A-tRNA^Gln^ interaction surface to a near amino acid level. To better understand binding of hTRMT2A to tRNA, we used the AlphaFold homology model of hTRMT2A (47), removed protein areas with low confidence prediction scores (< 70 pLDDT, https://alphafold.ebi.ac.uk/entry/Q8IZ69) and mapped the identified crosslinks onto the model. Superposition of this homology model to the crystal structure of TrmA bound to a tRNA T-Loop allowed us to confidently superpose a full tRNA^Phe^ (PDB ID: 4TRA) into the homology model (**Figure 8**). The derived hTRMT2A-tRNA^Phe^ model is consistent with experimental observations presented. For example, all three hTRMT2A domains – RBD, Central and methyltransferase domain – are in close contact with the modeled tRNA. Hence, this multi-domain interaction of full tRNA is in full agreement with our EMSA experiments, where tRNA binding by hTRMT2A FL was much stronger than binding by its individual subdomains (**Figure 2**). In addition, observed crosslinks are localized around the catalytic center of the model (**Figure 3, Supplementary Figure 3**) and map to the region into which the tRNA T-Loop is predicted to be inserted for methylation (19). Importantly, the crosslink of amino acid C463, found in all four datasets, resides near the modeled tRNA T-Loop stem. In our homology model of hTRMT2A the crosslinked C270 is placed between T-Loop and D-Loop of the tRNA to potentially interrupt tertiary interactions and render U54 accessible for methylation, as reported for *E. coli* TrmA (15). Both amino acids are also found crosslinked to a dinucleotide UU, which suggests their proximity to the T-Loop U54-U55 or the D-Loop U18-U19 of tRNA. The crosslink of residue H386, found in three of four datasets, is positioned close to the catalytic center, but further away from the modeled tRNA (**Figure 8A**). Several more residues found in less than 3 datasets such as Y331, W176, and C486 positioned close to the catalytic center can be explained with our model. Other crosslinks would require structural rearrangements to be close enough to the RNA such as C191, C225, C329, K559, C551, and C225. As the RBD is connected via an unstructured linker (aa 148-178) to the methyltransferase domain, positioning the RBD with respect to the RNA bears considerable uncertainty (**Supplementary Figure 14**). However, our cross-linking data indicate that the RBD binds RNA with its ß-sheet surface as well as with the flexible loop regions close to crosslinked H83. To account for these experimental data, the position of the RBD was adjusted manually with respect to tRNA (**Supplementary Figure 14, zoom**). The manually curated orientation of the RBD appears to be suitable to form the crosslinking-data suggested interface and thus stabilize the hTRMT2A-tRNA complex.

**Figure 8.**
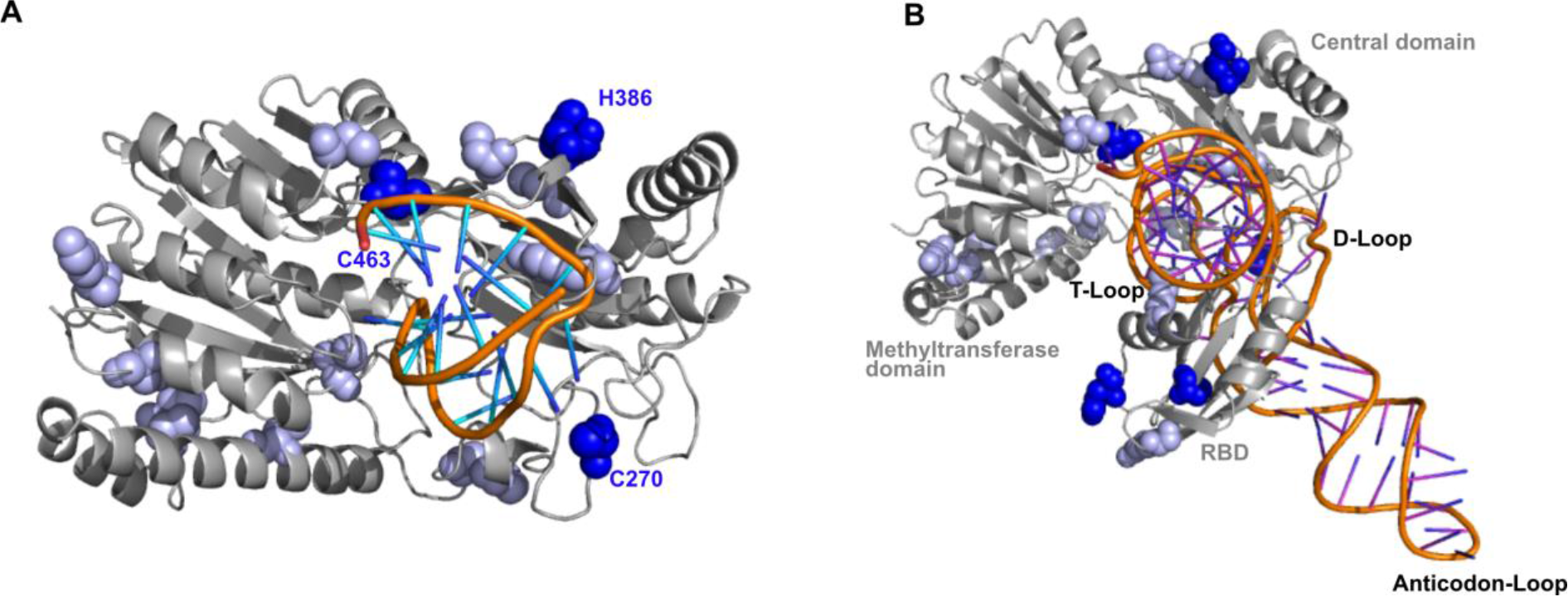
Model of hTRMT2A-tRNA complex. A. hTRMT2A AlphaFold homology model (grey) overlayed with the T-Loop from *E. coli* TrmA structure (PDB ID: 3BT7; orange). Consistently (blue spheres, 3 of 4 datasets) and less consistently (bright blue spheres, < 3 of 4 datasets) crosslinked amino acids were mapped onto the model. **B** Hybrid model consisting of the hTRMT2A AlphaFold homology model (grey) and an overlayed experimental full tRNA structure (PDB ID: 4TRA) based on the superposition with the *E. coli* TrmA T-Loop co-structure (PDB ID: 3BT7). Crosslinked amino acids from mass spectrometry experiments are highlighted (coloring as in A) and mapped onto the hybrid model. Models were generated with PyMOL.

Interestingly, our model is also consistent with hTRMT2A’s limited substrate selectivity with respect to RNA binding and methylation, because large proportions of the tRNA such as the anticodon stem and loop are not bound by hTRMT2A. Deviations in these structures might be tolerated, which would explain the binding of other RNAs than substrate tRNA such as stem- looped structured RNA or mRNA (**Figure 1**, **Figure 8**, (20)).

Despite the very consistent insights that arise from combining our experimental data with an advanced homology model, a full comprehension of the hTRMT2A-tRNA interface would require an experimentally determined high-resolution co-structure.

### The hTRMT2A interactome and future implications

With BioID and Co-IP we combined the strength of both methods to dissect the hTRMT2A interactome. BioID is superior in recapitulating the physiological nano-scale environment of hTRMT2A, as biotinylation of proteins occurs within intact cells. Moreover, transient interactions can be captured. However, BioID does not allow to distinguish between direct interactors and proximal, non-binding proteins for instance in a very crowded environment such as the nucleolus. In contrast, Co-IP confidently identifies proteins that are part of the same co-complexes including direct and indirect interactions of the bait protein. Moreover, Co-IP bears the risk of detecting unphysiological interactions that occur only after cell lysis. For RBPs with lower specificity, in particular, the risk of false positives is high. Having these differences in mind, we propose that most confident protein interactors can be identified by merging Co-IP and BioID data. A total of thirteen proteins were found both in BioID and Co-IP datasets from hTRMT2A WT cell lines: NCL, TRIM26, hTRMT2A, SSB, TARBP2, LARP7, PPM1G, UPF1, IGF2BP1, NPM1, SET, SPINDOC and RPL4. Some hits (SSB, TARBP2) were previously identified in orthogonal studies, which may serve as positive controls and indicate that our combinatory interactome study is robust and reliable (48). Proteins found in BioID and Co-IP from both hTRMT2A WT and CDM cell lines were SSB, TARBP2 and SPINDOC, which we propose as most likely protein interaction candidates independent of its enzymatic activity. Together, these cross-validated candidates indicate biological pathways hTRMT2A likely contributes to. For example, SSB acts as 3’ poly(U) terminus dependent transcription factor of RNA polymerase III-dependent transcripts by facilitating proper folding and maturation of target RNA. The best-studied function of SSB is its association with the pre-tRNA 3’ trailer, which supports its proper endonucleolytic cleavage (49). In yeast, early occurring tRNA modifications were installed while the pre-tRNA was still associated with the *S. cerevisiae* homolog of SSB (50, 51). Therefore, hTRMT2A could indeed methylate U54 of pre-tRNA, while the RNA is still associated with SSB. In that context, hTRMT2A and SSB might interact via bound tRNA as inferred from our BioID/Co-IP data. Also, RISC loading complex subunit TARBP2 was highly enriched both in BioID and Co-IP datasets. Interestingly, SSB prevents mischanneling of tRNA fragments into the human microRNA pathway, which might establish a further link between hTRMT2A, SSB and TARBP2 (52).

Lastly, hTRMT2A might be involved in rRNA processing, as GO-terms for “rRNA processing” and “ribosome biogenesis” were enriched. Related interaction candidates were identified among the top BioID and Co-IP hits (**Figure 5**) such as Nucleolin (NCL), which is involved in pre-rRNA transcription and maturation (53) or RPL4, which is part of rRNA processing as well as ribosome biogenesis pathways (54). These implications together with the fact that the human ortholog TRMT2B modifies both mt-tRNA and mt-rRNA, motivated us to assess the m^5^U status of cytosolic rRNA isolated from hTRMT2A WT and hTRMT2A KD. We observed that rRNA from hTRMT2A WT and KD showed only a minimal fraction of m^5^U modified uridines, which was similar to the background levels of modifications that were known to be absent in rRNA (43) (**Supplemental Figure 13C**). This suggests that m^5^U is not installed on rRNA in the TRMT2A WT and KD cell lines used in this study (**Figure 7B**). Therefore, we conclude that BioID/Co-IP identified proteins are enriched in these pathways not because hTRMT2A modifies rRNA directly, but potentially due to a non-enzymatic, secondary functions in ribosome/rRNA biogenesis.

As no impact of TRMT2A depletion on protein synthesis was observed (34), we aimed at addressing the question whether reduced hTRMT2A and therefore m^5^U levels in tRNA or other RNA species impact translation fidelity. Indeed, in a translation fidelity assay we observed increased translation errors in hTRMT2A KD cells over WT using the E145Q renilla variant, but not with a D120N renilla variant (**Figure 7C**). Differential translation error for one type of mutation but not another indicates a selective loss of fidelity. One possible explanation is a differential bias in tRNA codon abundance of glutamic and aspartic acid in HEK 293t cells (55). In this context, it is worth noting that hTRMT2A has been identified in a *Drosophila* enhancer/suppressor screen as poly-glutamine (polyQ) disease modifier. Reduced levels of hTRMT2A were associated with decreased polyQ aggregates and photoreceptor death (56). While the rescue effect of decreased hTRMT2A levels on polyQ aggregate formation could be recently validated in human cells, the mode of action has remained elusive (34). Our observation here suggests that increased translation error rates might cause a stochastic exchange of glutamine for instance against glutamate and therefore disruption of long neurotoxic polyQ glutamine (polyQ) stretches. The tremendous impact of polyQ-stretch disruption on polyQ aggregation has been shown earlier (57). Importantly, although our data point to a contribution of tRNA methylation it remains to be proven whether an increase in translation error rate is indeed due to hypomethylation of tRNA, mRNA, or other cellular processes such as tRF generation (20, 24, 45). As the reduction in translation fidelity is only modest and possibly restricted to certain amino acids, it is not surprising that hTRMT2A WT and KD do not show any apparent phenotypic differences in culture. This is in line with observations in yeast, where depletion of the ortholog Trm2 was not correlated with an adverse phenotype (9).

Together our *in vitro* RNA binding and methylation data and protein-interactome studies contribute to a basic understanding of the underexplored hTRMT2A molecular and cellular function. Next to its major role in tRNA methylation, these data also uncovered another role of hTRMT2A in protein translation fidelity via yet to be determined mechanisms. Furthermore, since TRMT2A has been described as a modulator of neurotoxicity in poly-glutamine aggregation disorders (34, 56), this study might help to establish hTRMT2A as a target for therapeutic intervention.

## Acknowledgments

We thank the Core Facility Metabolomics and Proteomics (CF-MPC), especially Ann-Christine König for providing expertise, data recording, and analyses. Further, we thank Monika Raabe for help with crosslinking and MS, Sebastian Iben for help setting up the translation fidelity assay, and Markus Müller for help with MS analyses of rRNA modifications. We thank Irmgard Diepolder and Vera Roman for technical support.

