## Supplementary Figures 1-14 for "Human TRMT2A methylates tRNA and contributes to translation fidelity"

1 Institute of Structural Biology, Helmholtz Zentrum München - German Research Center for Environmental Health, Neuherberg, Germany.
2 Institute of Pharmaceutical Biotechnology, Ulm University, Ulm, Germany.
3 Bioanalytical Mass Spectrometry Group, Max Planck Institute for Multidisciplinary Sciences, Göttingen, Germany.
4 Institute of Clinical Chemistry, University Medical Center Göttingen, Göttingen, Germany.
5 Department of Chemistry and Biochemistry, Ludwig-Maximilians University Munich, München, Germany.
6 Metabolomics and Proteomics Core, Research Unit Protein Science, Helmholtz Zentrum München - German Research Center for Environmental Health, Neuherberg, Germany.
7 Department of Neurology, Faculty of Medicine, RWTH Aachen, Aachen, Germany.
* Correspondence should be addressed to:
Aaron Voigt, Dierk Niessing

### Supplemental Figures

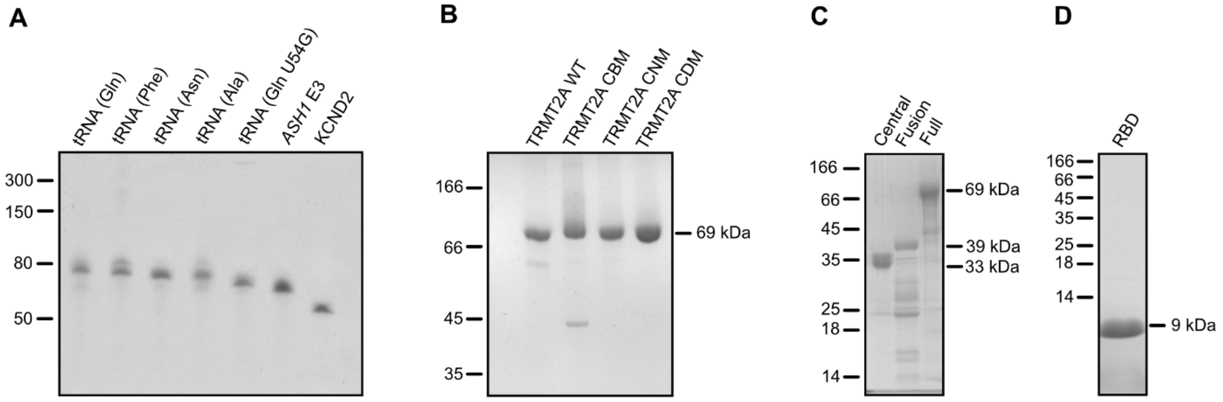


**Supplementary Figure 1 – Purity of recombinant protein and *in vitro* transcribed RNA for EMSAs and MTase Glo assays (Figure 1, 2, 5).**

**A** *In vitro* transcribed, PAGE purified RNA (250 ng) was separated on an 8 % denaturing PAGE, stained with ethidium-bromide. Staining displays high RNA purity. **B** hTRMT2A WT and mutants (10 µM) were run on 12.5 % SDS PAGE, stained with Coomassie. Protein preparations show high purity. **C** and **D** display recombinantly purified hTRMT2A protein domains separated on 12 % (C) and 19 % (D) SDS PAGE and stained with Coomassie.

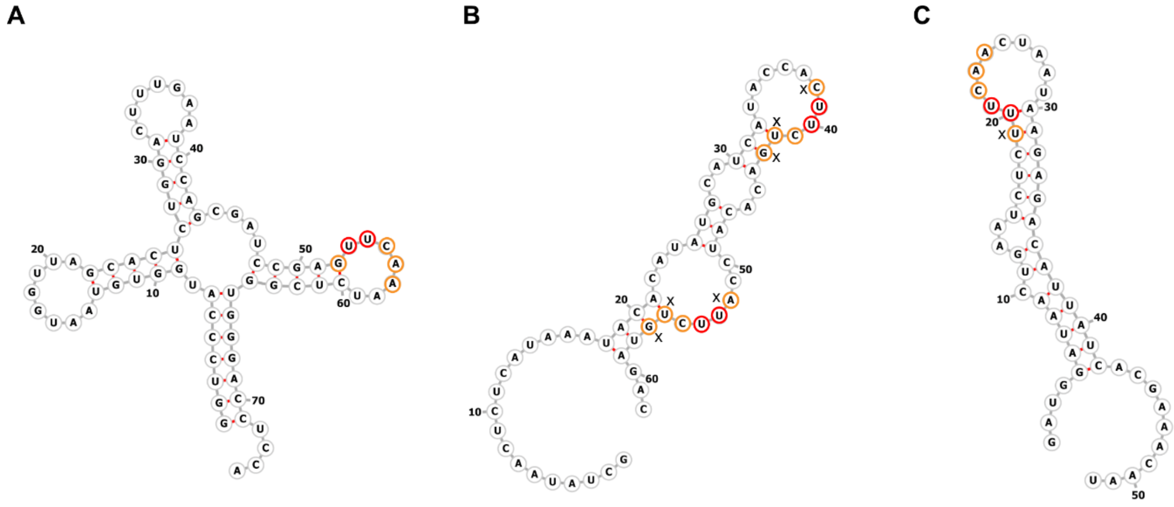


**Supplementary Figure 2 – Predicted RNA structures**

**A** Predicted RNA fold of tRNA$^{Gln}$ (75 nt). **B** Predicted RNA fold of KCND2 mRNA, a potential hTRMT2A RNA target (62 nt). **C** Predicted RNA fold of the *E3* element of *S. cerevisiae ASH1* mRNA (51 nt). Target uridine and adjacent uridine are highlighted in red. Other residues, which are predicted to be part of the hTRMT2A sequence motif are highlighted in orange. RNA folds were predicted with the RNAfold Webserver (1).

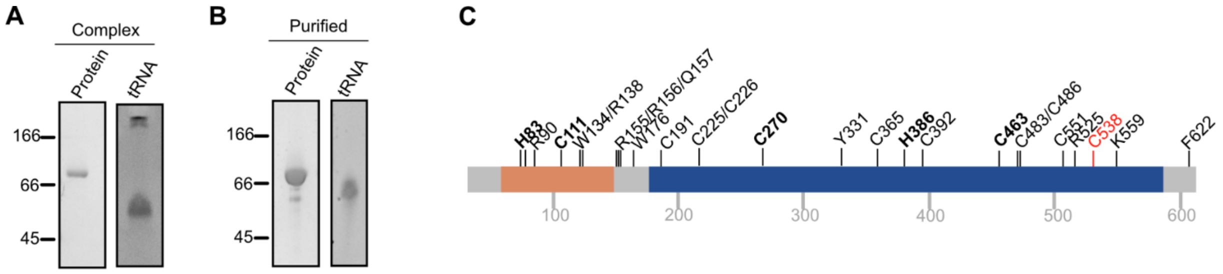


**Supplementary Figure 3** – **Purification and crosslinking results of the hTRMT2A-tRNA$^{Gln}$ complex.**
**A** SDS PAGE (12 %) and denaturing PAGE (8 %) of recombinantly purified hTRMT2A FL (1-625) and *in vitro* transcribed tRNA$^{Gln}$, respectively, as used for complex reconstitution experiments. RNA was stained with ethidium-bromide and protein with Coomassie. **B** Protein-tRNA complex as purified with size exclusion chromatography. Samples were separated on SDS PAGE (12 %) for protein and denaturing PAGE (8 %) for RNA visualization. **C** Results of crosslinking experiments mapped on a linear representation of hTRMT2A. Crosslinked amino acids in three or four of four experiments (bold letter), conserved catalytic residue C538 (red) are highlighted. Crosslinked amino acids in less than three of four experiment are not highlighted**.** Crosslinked amino acids are distributed across the RBD (orange) and the methyltransferase domain (dark blue).

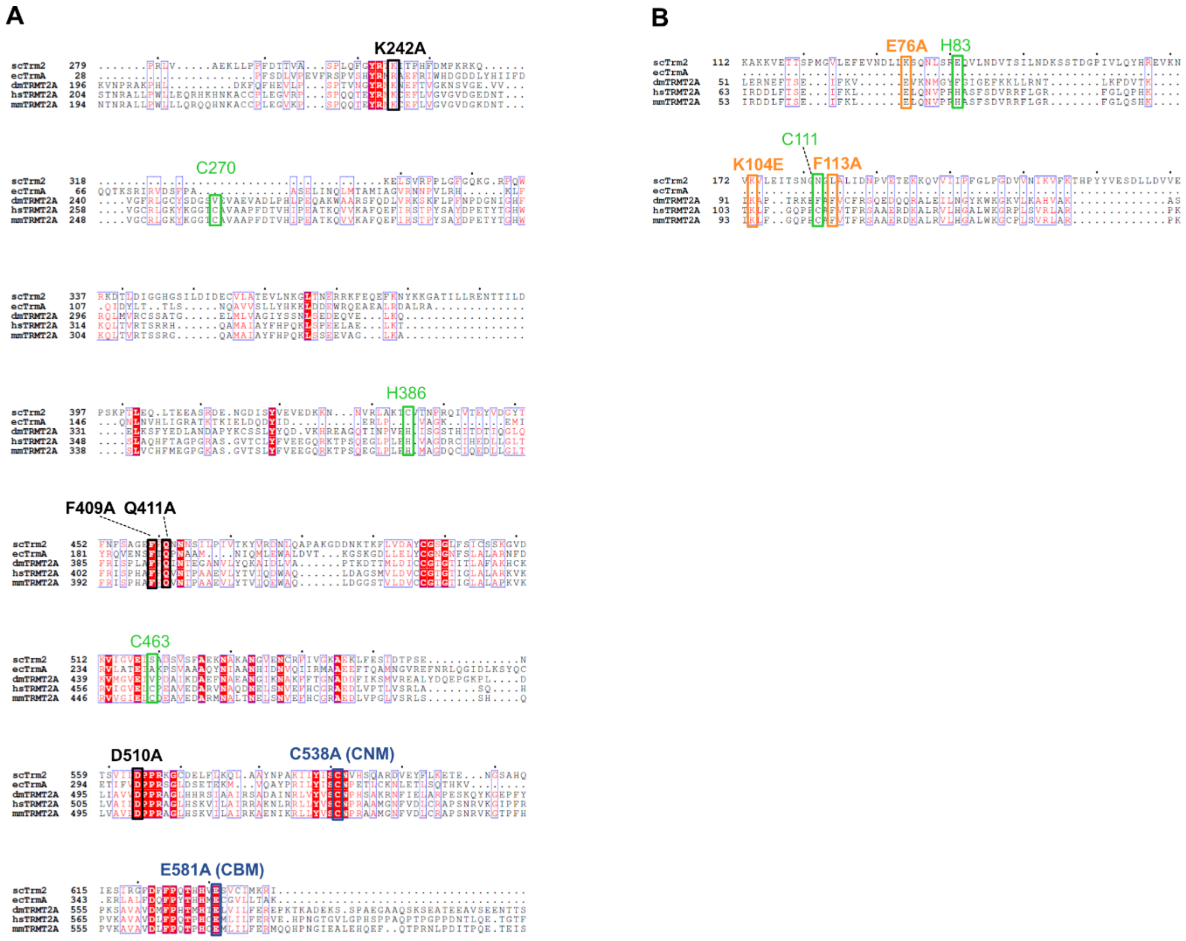


**Supplementary Figure 4 – Multiple sequence alignment of *Escherichia coli* (ec), *Saccharomyces cerevisiae* (sc), *Drosophila melanogaster* (dm), *Mus musculus* (mm), homologs of *Homo sapiens* (hs) TRMT2A.**
**A** Multiple sequence alignment of the methyltransferase domain. CBM (blue), CNM (blue) and CDM (black) mutations are labelled. C538A mutation occurs in the CNM and CDM mutant and is one of the crosslinked amino acids. Amino acids crosslinked in three or four of four experiments are shown (green). **B** Multiple sequence alignment of RBD. CDM mutations (orange) and crosslinked amino acids (green) are shown. ClustalW algorithm was used for multiple sequence alignment. Highly conserved amino acids are highlighted in red. Visualization of alignment was done with ESPript,version 3.0 (2).

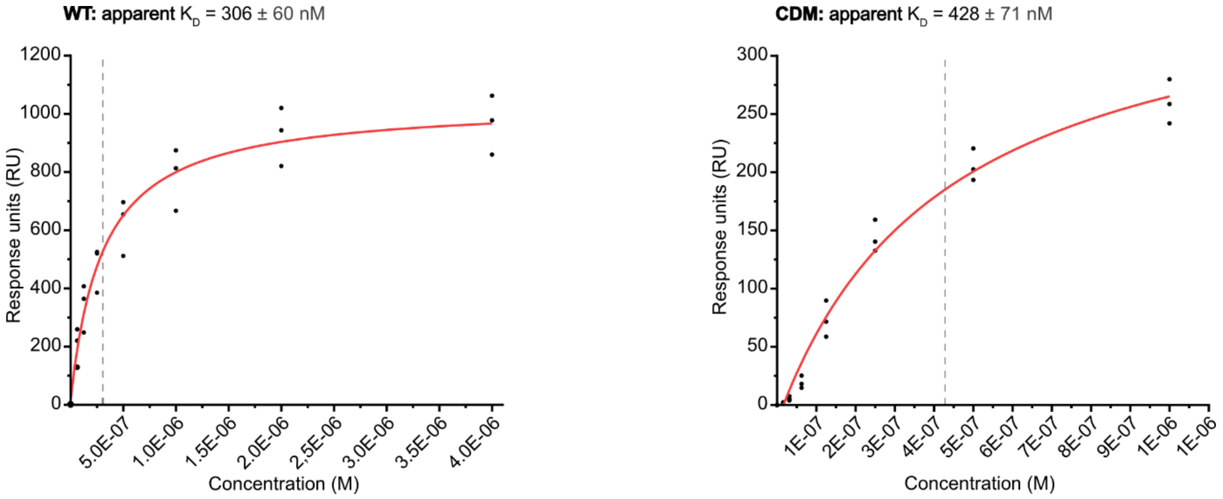


**Supplementary Figure 5 –K$_D$ fits of hTRMT2A WT and CDM SPR experiments.**
For K$_D$ curves protein concentrations were plotted against response units and fitted with a steady-state affinity model to derive apparent K$_D$ values. SPR experiments were performed with biotinylated tRNA$^{Gln}$ coupled to streptavidin-coated surface of a SA-chip. Injection series of 62.5 – 4000 nM (WT) and 15.6 – 1000 nM (CDM) resulted in shown K$_D$ curves and apparent K$_D$ values (dashed line) of 306 ± 60 (WT) and 428 ± 71 (CDM), respectively. Fits of triplicate measurements are shown.

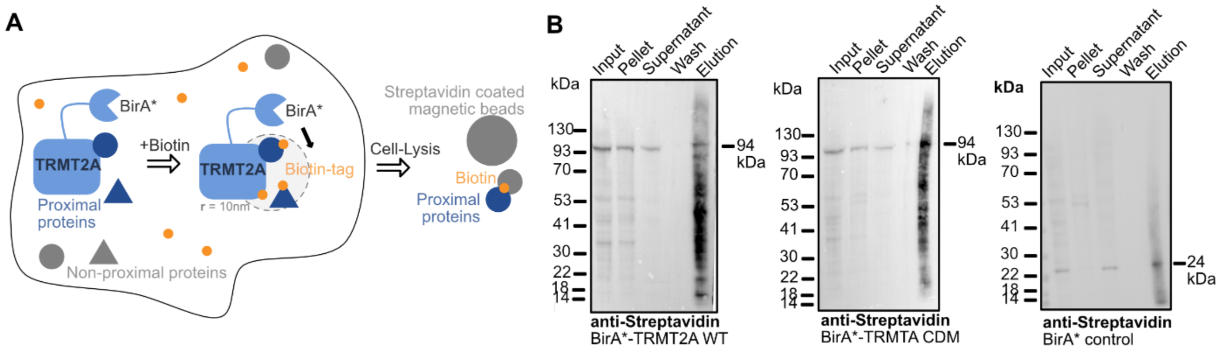


**Supplementary Figure 6 – BioID experiment**
**A** Scheme of a BioID experiment. Fused BirA* ligase biotinylates proximal proteins within a radius of 10 nm. Biotinylated proteins were captured with Streptavidin-coated beads and subjected to mass spectrometry analyses. **B** Western blot of representative replicates from BirA* control, BirA*-hTRMT2A WT and BirA*-hTRMT2A CDM. The main band of the protein of interest in the input sample runs at the correct height. The wash fractions were empty, which indicates sufficient washing. Elution sample illustrated several captured biotinylated proteins over a large range of protein sizes. Primary antibody used: HRP-conjugated anti-Streptavidin (1:10.000, Thermo, #N100).

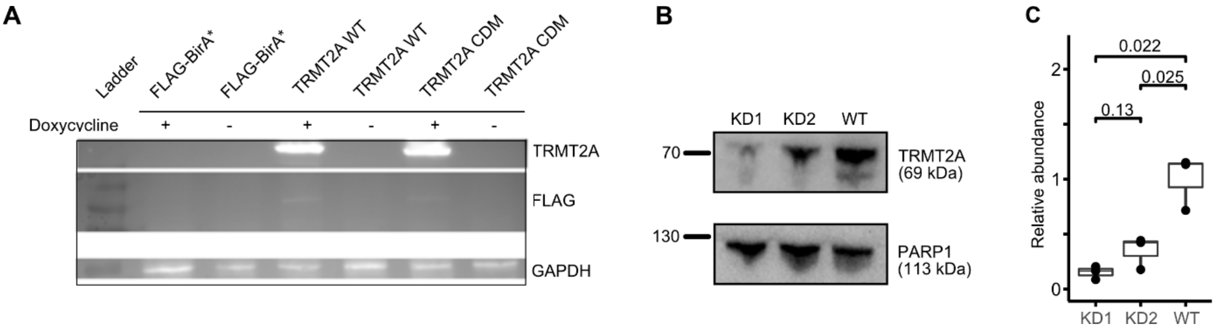


**Supplementary Figure 7 – Western blot of BioID cell lines and hTRMT2A KD cell lines**
**A** Western blot of FLAG-BirA* control cell line, as well as FLAG-BirA* tagged hTRMT2A WT and hTRMT2A CDM overexpression cell lines used in BioID experiments. Upon supplementation of 1:1000 doxycycline, overexpression of hTRMT2A is apparent in hTRMT2A WT and CDM cell lines, but not in the BirA* control. **B** Stable RNAi-mediated knockdown of hTRMT2A (KD1, KD2) in HEK 293t cells was achieved with shRNA lentiviral transduction particles Representative Western blot of hTRMT2A knockdown cell lines, as well as hTRMT2A WT control. A clear reduction of hTRMT2A levels is visible in both knockdown cell lines compared to WT, with a stronger reduction in KD1. **C** Box plot of the densitometric quantification of hTRMT2A levels in KD1, KD2 and WT cell lines using three independent replicates show significant reduction of hTRMT2A levels in KD1 ($p$ = 0.022) and KD2 ($p$ = 0.025). $p$ values were calculated using an unpaired two-sided Student´s $t$-test. Primary antibodies used: rabbit anti-TRMT2A (1:1000; Abcam, ab205616), rabbit anti-PARP1 (1:1000; Sigma-Aldrich, HPA045168), mouse anti-FLAG (1:2000; Sigma, #F3165), mouse anti-GAPDH (1:500, DSHB-hGAPDH, #2G7). HRP-conjugated secondary antibody used: goat anti-rabbit (1:10,000; Abcam, ab6721), sheep anti-mouse (1:10.000; GE Healthcare, #NXA931V), donkey anti-rabbit (1:10,000; GE Healthcare, #NA934V).

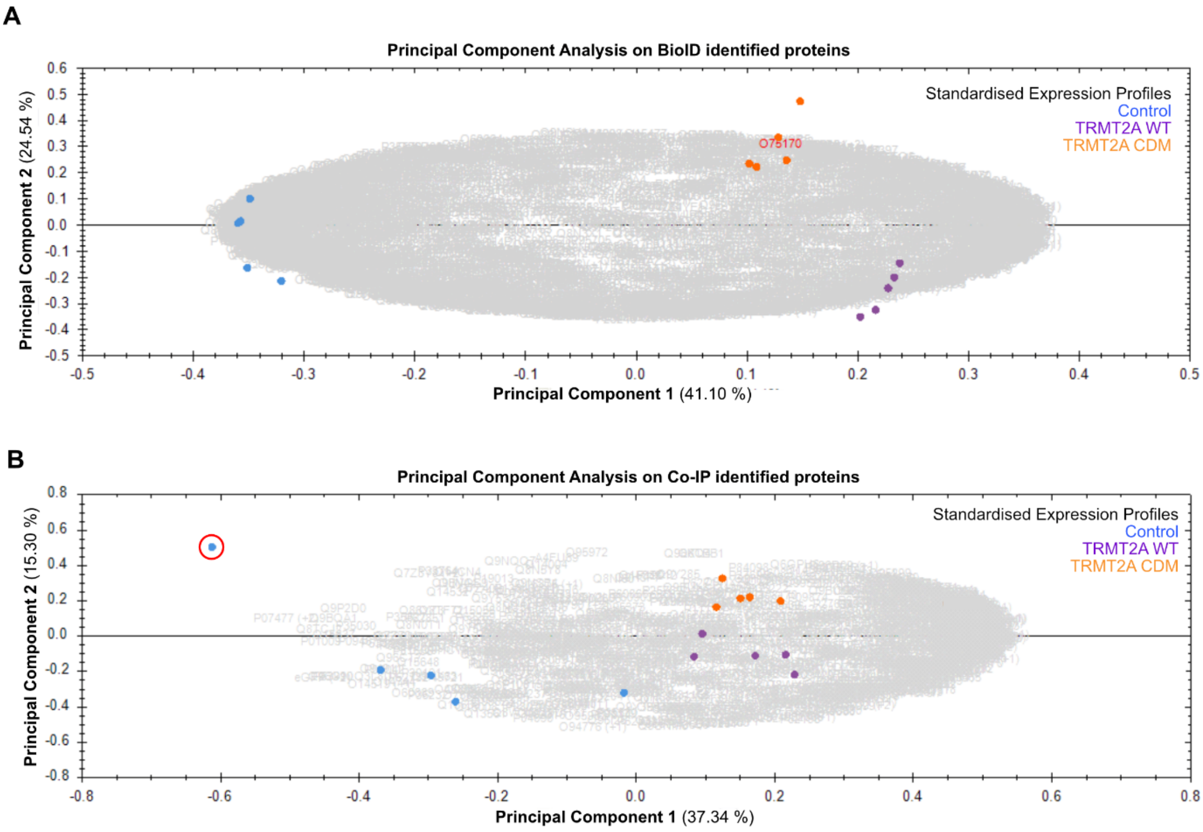


**Supplementary Figure 8 – Principal component analysis BioID and Co-IP data**
**A** PCA on BioID identified proteins with quintuplicates of BirA* control (blue), BirA*-hTRMT2A WT (violet) and BirA*-hTRMT2A CDM (orange) shows significant differences between samples from different conditions. **B** PCA on Co-IP identified proteins with quintuplicates of FLAG control (blue), FLAG-hTRMT2A WT (violet) and FLAG-hTRMT2A CDM (orange) shows significant differences between samples from different conditions. Outlier from control sample is encircled in red. PCA analysis was performed with Progenesis QI software and uses feature abundance levels across runs to determine the principal axis of abundance variation.

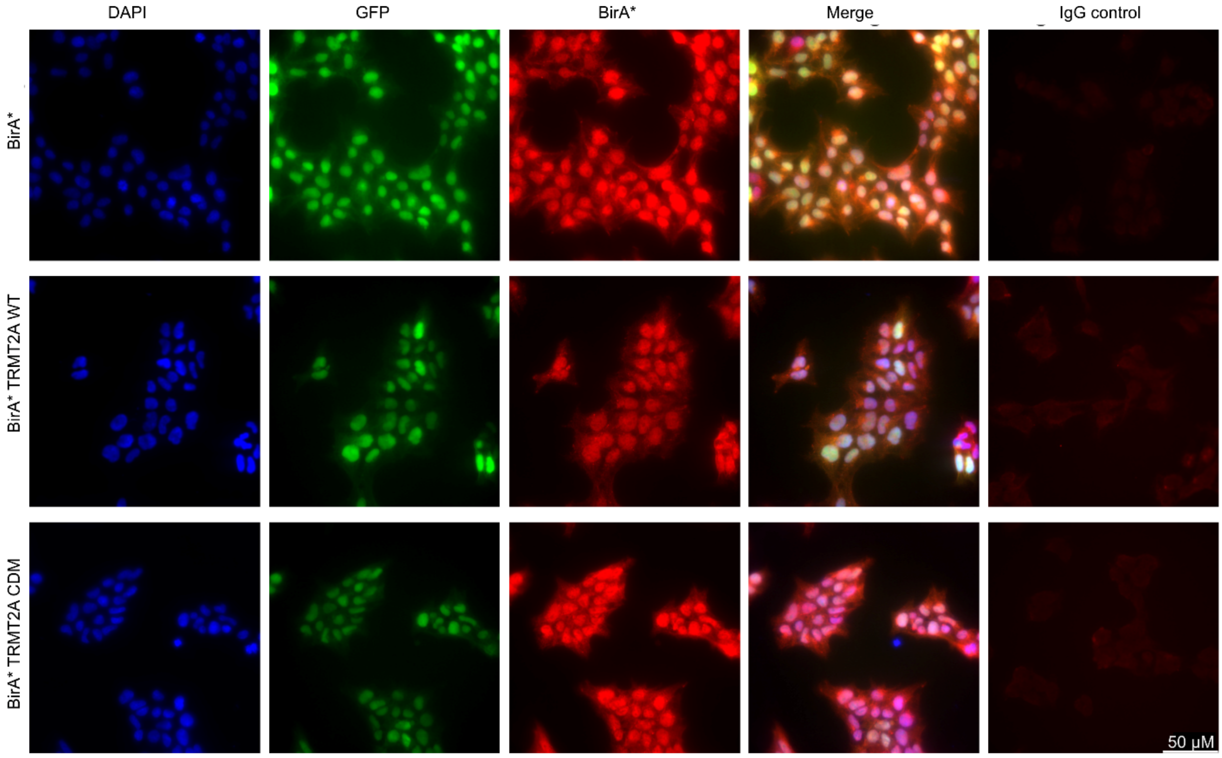


**Supplementary Figure 9 – Immunostainings of BirA* cell lines (Control, hTRMT2A WT and CDM)**
Images of HEK 293t cells stably expressing the FLAG-BirA* control, FLAG-BirA* hTRMT2A WT and FLAG-BirA* hTRMT2A CDM. The fusion protein is predominantly localized to the nucleus, but also to the cytoplasm. GFP serves as expression control, IgG staining as background control and DAPI as nuclear stain. Primary antibodies used: mouse anti-BirA (1:10, Novus Biologicals, #5B11c3-3). Secondary antibody used: goat anti-Mouse IgG Alexa Fluor 647 (1:2000, Invitrogen A-21235).

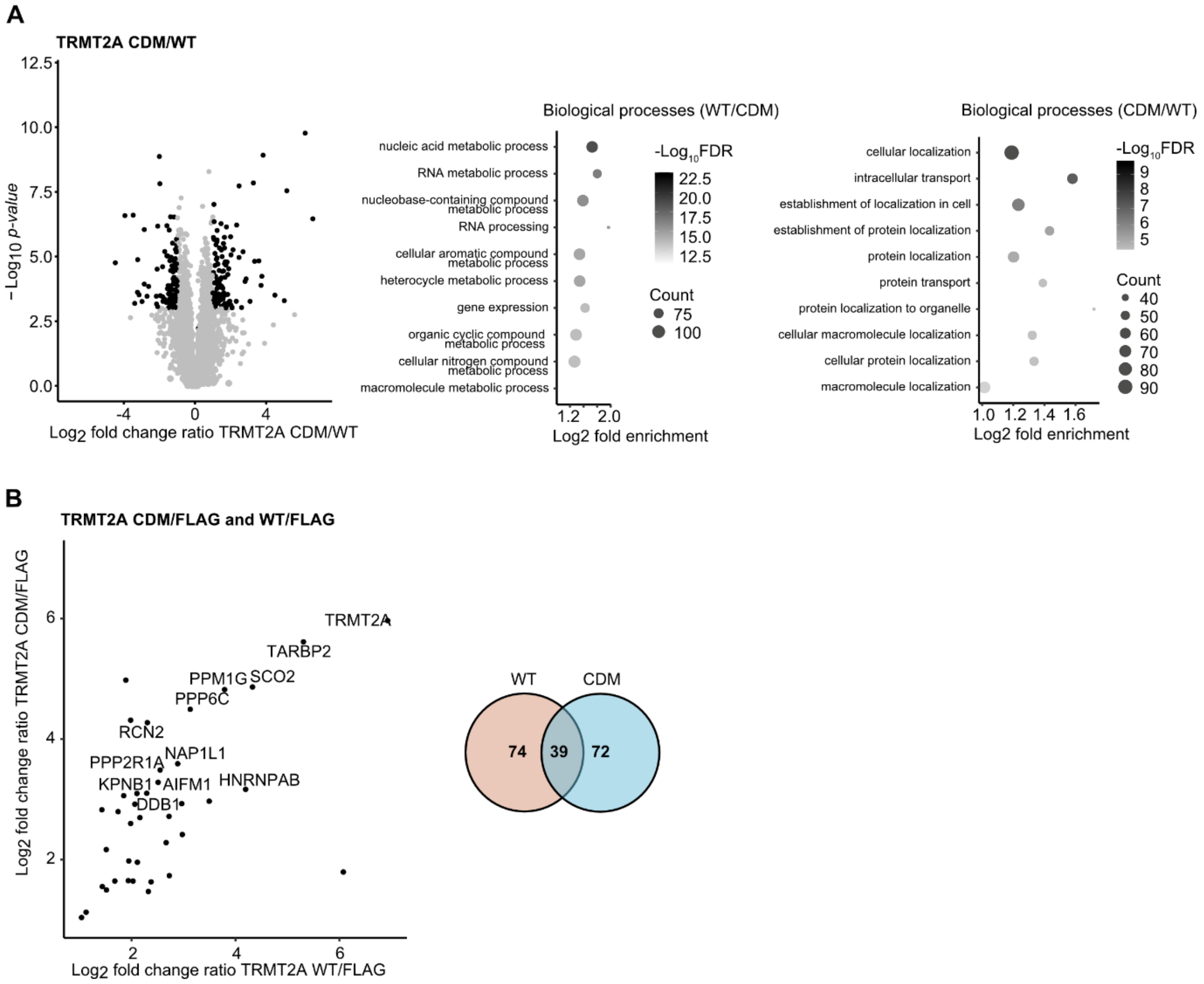


**Supplementary Figure 10 - Volcano plot of BioID experiment and Co-IP overlap**
**A** Volcano plot hTRMT2A CDM over hTRMT2A WT from BioID experiment shows similar fold changes of enriched proteins. Log2 fold change ratio of TRMT2A CDM/WT was plotted against Log10 p-value. Ratio cut-offs were > 2 and < 0.5 and significance cut-off, with a p-value < 0.05. Hits in agreement with these thresholds are highlighted black. Top 10 most enriched GO-terms for biological processes and cellular compartments of hTRMT2A WT/CDM and hTRMT2A CDM/WT ratios show depletion of RNA associated processes in the hTRMT2A CDM mutant. GO-terms were ranked according to false discovery rate (FDR) as computed with the fisher's exact test. Graphs illustrate GO-terms plotted against Log2 fold enrichment. Node color represents -$Log_{10}$FDR. Node size represents counts per GO-term. GO-terms were visualized in R. **B** Plot shows the hTRMT2A WT/FLAG and hTRMT2A CDM/FLAG ratios. Venn diagram depicts total enrichment of 74 proteins in WT (orange) and 72 in CDM dataset (blue) with an overlap of 39 proteins.

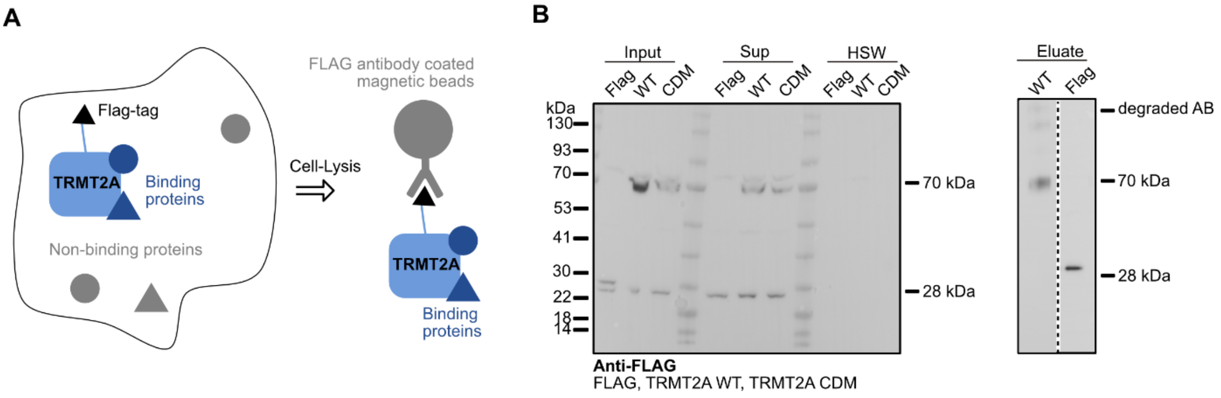


**Supplementary Figure 11 – Co-IP experiment**

**A** Principle of a Co-IP experiment. Fused FLAG peptide allows for enrichment of hTRMT2A and all bound proteins using FLAG antibody coated beads. Bound proteins were identified with mass-spectrometry. **B** Western blot of representative replicates from FLAG control, FLAG-TRMT2A WT and FLAG-TRMT2A CDM. The main band of the protein of interest in the input sample runs at the correct height. The high salt wash fractions (HSW) were empty in line with sufficient washing. Elution samples from FLAG-control and FLAG-TRMT2A WT illustrate successful enrichment of protein of interest at the correct height. Primary antibody used: rat anti-FLAG (1:10; in-house, IgG1, monoclonal) Secondary antibody used: HRP-conjugated mouse anti-rat (1:1000; in-house, polyclonal).

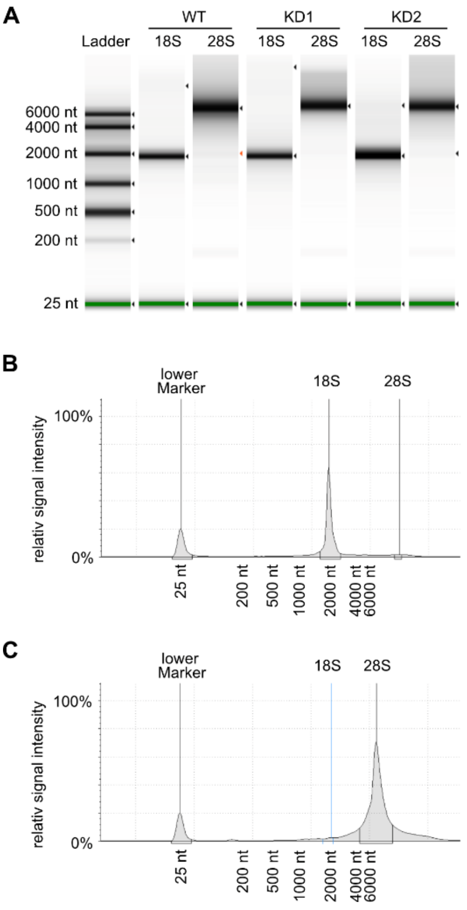


**Supplementary Figure 12 – Agilent Tape station analysis of purified rRNA**

**A** Representative Agilent Tape station measurement of rRNA from hTRMT2A WT, KD1 and KD2 cell lines shows high purity and correct size. **B** Electropherogram of 18S rRNA indicates effective separation from 28S rRNA and no tRNA contaminations. **C** Electropherogram of 28S rRNA indicates effective separation from 18S rRNA and no tRNA contaminations. Measurements were performed according to manufacturer's recommendations.

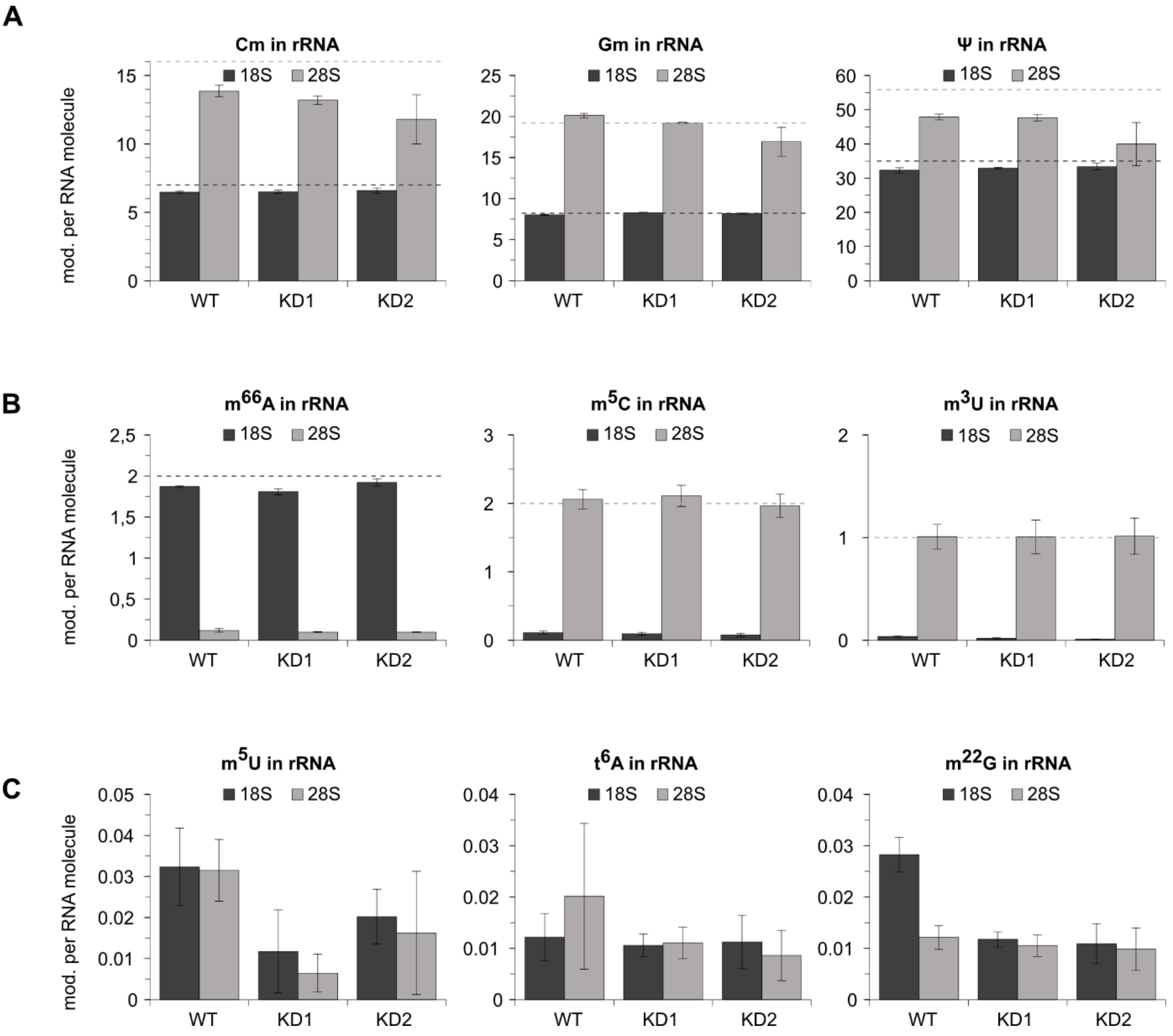


**Supplementary Figure 13 – Mass spectrometry analysis of modifications in 18S and 28S rRNA**

**A** LC-MS/QQQ analysis of nucleosides of 18S and 28S rRNA isolated from hTRMT2A WT, KD1 and KD2 cell lines. Number of modifications per RNA molecule for Cm, Gm and ¤ modifications in rRNA shows good agreement with reported numbers for 18S and 28S rRNA (dashed lines, Modomics). **B** Exemplary data of modifications that are only found in rRNA from one subunit; $m^{66}A$ occurs in 18S rRNA, and $m^5C/m^3U$ in 28S rRNA. **C** Exemplary data of modifications that were not found in rRNA previously; $m^{22}G$, $t^6A$ and prospectively $m^5U$. Quantitative mass spectrometry measurements have been performed in biological triplicates on an Agilent 1290 Infinity equipped with a variable wavelength detector (VWD) combined with an Agilent Technologies G6490 Triple Quad LC/MS system with electrospray ionization.

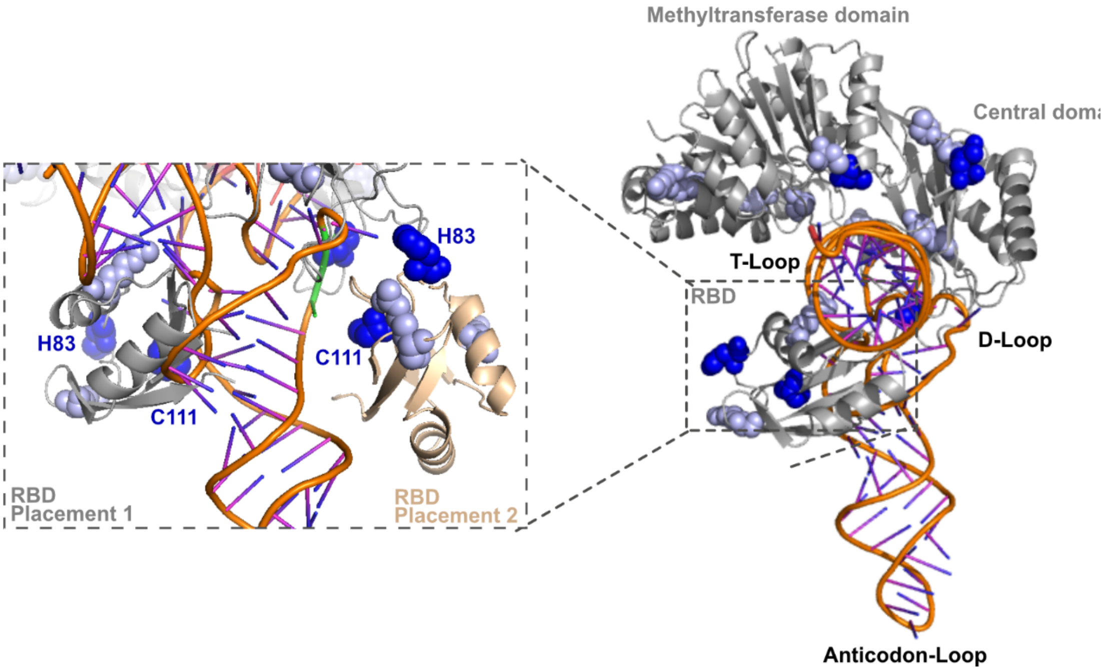


**Supplemental Figure 14 - Hybrid model of hTRMT2A-tRNA binding and RBD position adjustment**
Hybrid model consisting of the hTRMT2A homology model (AlphaFold) and full tRNA structure (right hand side) positioned in a similar orientation as in Figure 8B. The zoomed insert (left hand side) shows a 90 ° rotated view of the model. The original RBD placement 1 (grey) arises from the rigid body overlay of the hTRMT2A homology model with the *E. coli* TrmA T-Loop co-structure (PDB ID: 3BT7) and subsequent superposition of a full tRNA$^{Phe}$ (PDB ID: 4TRA) to the T-Loop. Using the consistent crosslinks between the RBD and the tRNA an adjusted RBD placement 2 (beige) is proposed, which brings the crosslinked residues H83 and C111 in close proximity to the tRNA.
